# Developing a multi-modal neuroimaging-based BrainAge model across childhood

**DOI:** 10.64898/2026.05.19.725847

**Authors:** Shi Yu Chan, Pei Huang, Ai Ling Teh, Arshia Naaz, Jasmine S.M. Chuah, Zhen Ming Ngoh, Janice Lee, Aisleen M.A. Manahan, Xavier Y.H. Lim, Marielle V. Fortier, Helen J. Zhou, B.T. Thomas Yeo, Yap-Seng Chong, Peter Gluckman, Johan Eriksson, Rajkumar Dorajoo, Dennis Wang, Michael J. Meaney, Ai Peng Tan

## Abstract

BrainAge models hold promise as a clinical biomarker for developmental brain health, especially in childhood when there is the potential for early intervention. To distinguish between normative developmental variance and pathological divergence, BrainAge models should reflect the dynamic and diverse neurodevelopmental processes that occur in distinct developmental windows across childhood. We utilized multi-modal neuroimaging data from three pediatric cohorts covering ages 4 to 13 years (n = 1005, 2126 scans), split into Train and Test datasets. Twelve sex-stratified BrainAge models were built stratified by type and different combinations of neuroimaging features. Model types were “Full-Span” models covering the full age range, and “Phase-Specific” models split into early- and late-childhood. We first compared BrainAge estimates in the Test dataset amongst our candidate models, then benchmarked the best-performing model against published pre-trained models and DNA-based biological age measures. Our findings show that a BrainAge model that was phase-specific and consisted of both structural and functional features (cortical thickness, subcortical volumes, and functional network integration measures) showed good prediction of age and best distinguished between healthy and symptomatic subgroups. We present a proof-of-concept for developmental models supporting building BrainAge models of higher temporal resolution that align to different childhood developmental phases.

## 1. Introduction

The dynamic neurobiological processes that underlie childhood neurodevelopment are foundational to brain health and serve as predictors of later-life outcomes^1,2^. BrainAge models capture variations in these processes at the individual level, distilling complex, non-linear neuroimaging patterns into a singular multivariate index of neurobiological maturity^3^. The BrainAge literature shows success in aging and clinical cohorts as a predictive biomarker capable of identifying accelerated biological aging in individuals with Alzheimer’s Disease, Major Depressive Disorder, and Schizophrenia^4–10^. The quantified BrainAge gap showed associations with cognitive decline or illness severity^5,6,9^. The utility of BrainAge as a biomarker of developmental brain health depends on its ability to distinguish between normative developmental variance and pathological divergence. Optimizing these models enables earlier identification of at-risk individuals, allowing for intervention before functional or psychiatric symptoms manifest.

Widely-used BrainAge models in the literature are typically trained over large age ranges and are mostly based on anatomical data (volume, cortical thickness)^7,9,11–13^. With the challenges in collecting neuroimaging data in children, pediatric neuroimaging contributes only a small portion of the data. Hence, existing models are not designed to address neurodevelopment across childhood. Addressing this knowledge gap is of particular importance for the prediction of common mental health disorders, where an early age of onset necessitates early detection and intervention in childhood.

Integrating multi-modal neuroimaging data more effectively characterizes the diverse processes of childhood neurodevelopment than does single-modality approaches^14^. This integration is particularly vital given the spatiotemporal heterogeneity of brain maturation, where regional changes occur across distinct developmental windows^15^. While development occurs on a continuum, childhood can be partitioned into early-childhood (up to 8 years of age) and late-childhood (8–13 years) that reflect distinct neurobiological and functional epochs^15–20^. Early-childhood is characterized by the expansion of cortical grey matter and more local, uni-modal driven functional architecture^15,16^. The transition to late-childhood aligns with the peak of grey matter volume and the shift from a local to distributed functional organization with increasing long-range connections^15,17^, facilitating the emergence of higher-order reasoning and the development of social-evaluative processing^18,19^. Across multiple neuroimaging studies of structural and functional measures, changes in developmental trajectories occur within the age range of 6 to 11 years^16,20–23^. These shifts suggest that different neurodevelopmental processes define the early- and late-childhood phases.

We hypothesize that a BrainAge model that accurately captures the underlying neurodevelopmental changes across childhood will be sex-specific, include multi-modal neuroimaging data, and be specific to childhood developmental phases. To test this hypothesis, we used longitudinal multi-modal neuroimaging data from two pediatric cohorts in Singapore to build candidate BrainAge models for children from ages 4.0 to 13.5 years. We built sex-specific models (i) across the full age range (“Full-Span”) and (ii) split into two developmental phases (“Phase-Specific”) - early (ages 4 to 7.5 years) and late (ages 7.5 to 13.5 years) childhood. We also used different combinations of neuroimaging features. Three were uni-modal models consisting of either structural or functional data: cortical thickness (“S1”); cortical thickness with subcortical volumes (“S2”); functional network recruitment, integration and modularity (“F1”). Three were multi-modal models: modularity, functional recruitment and structural measures (“A1”); modularity, functional integration and structural measures (“A2”), and all features (“A3”). Sex-specificity in neurodevelopmental changes and their functional impact is well-established in the literature^22,24–26^ and was not specifically tested in this study.

We first compared BrainAge models across different features and type (Full-Span or Phase-Specific). BrainAge model performance was assessed in a Test dataset consisting of a held-out sample and an independent pediatric cohort enriched for learning difficulties. Model performance was assessed with median absolute error (MAE), BrainAge gap skewness, and the ability to distinguish between healthy and symptomatic subgroups. Second, we identified the most important neuroimaging features in the best performing candidate BrainAge model. Finally, we benchmarked our selected BrainAge model against pre-trained BrainAge models from the literature and against biological age measures.

## 2. Results

### 2.1 Cohorts and Data Split

Our data were drawn from the participants of three independent, Singaporean pediatric cohorts. The Growing Up in Singapore Towards healthy Outcomes (GUSTO) study^27,28^ and the Singapore Preconception Study of Long-Term Maternal and Child Outcomes (S-PRESTO) study^29^ are population-based birth cohorts involving multiple waves of data collection from mother-child dyads. Data collection time-points range from pregnancy to adolescence and include environmental exposures, behavioral measures and biological samples. In GUSTO, longitudinal neuroimaging data was collected at 5 time-points (Y4.5, Y6.0, Y7.5, Y10.5, and Y13.0 years of age; n = 671, 1792 scans). In S-PRESTO, neuroimaging data was collected once between ages 4 to 6 years (n = 120). The BRAin COnnectivity and Learning Difficulties (BRACO-LD) study is an ongoing pediatric symptomatic cohort. Children with learning difficulties were recruited through clinic referrals and community sources. Neuroimaging data was collected once between ages 4 to 8 years (n = 214). S-PRESTO and BRACO-LD were designed to have cross-cohort compatibility and pre-harmonized prior to data collection. Fig 1 depicts the data collection time-points. Detailed cohort characteristics are summarized in Supplement 1 with the study flow diagram and analysis numbers in Fig S1. Demographics and MRI quality check (QC) measures across the cohorts and time-points are summarized in Tables S1a-b.

**Figure 1:**
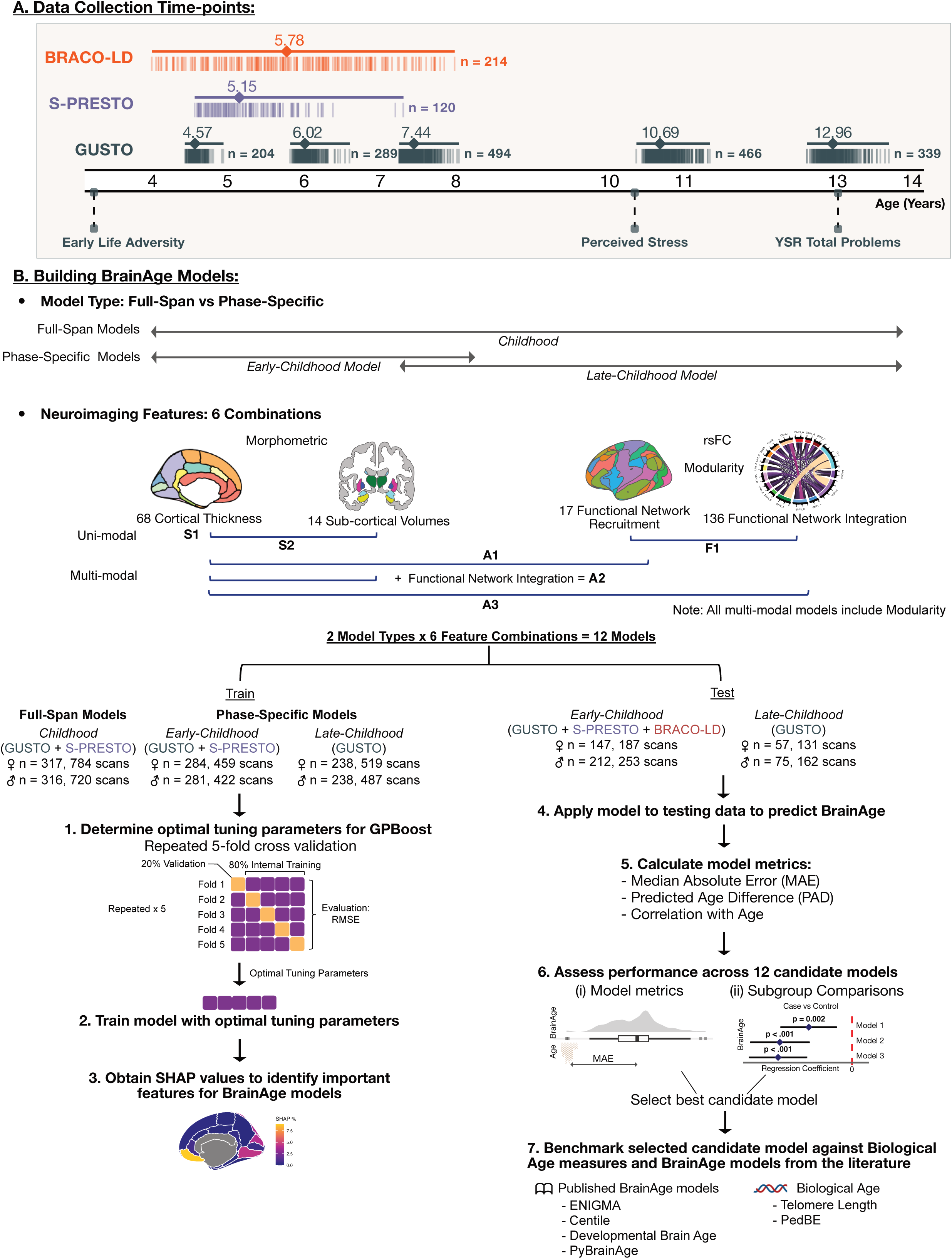
Data collection time-points and study design. (A) Neuroimaging data was collected from three pediatric cohorts from age 4 to 14 years, with repeated scans collected in the GUSTO cohort. Each vertical line represents an individual scan with the diamond representing the median age of the scan at each cohort/time-point. Early life environment variables and late childhood outcomes were also collected in GUSTO. (B) 12 candidate BrainAge models were developed based on different combinations of model type (Full-Span vs Phase-Specific models) and neuroimaging features. The full sample was split into Train and Test datasets, with BrainAge model training and feature importance determined in the Train dataset and model validation in the Test dataset. Predicted BrainAge from the best candidate model was then benchmarked against pre-trained BrainAge models from the literature and DNA-based biological age measures. Note: rsFC, resting state functional connectivity; RMSE, root mean square error; SHAP, Shapley Additive exPlanations; YSR, Youth Self Report

The total sample (n = 1005, 2126 scans) was split into a Train (n = 633, 1504 scans) and Test dataset (n = 372, 622 scans). Partitioning was done by participant ID to ensure that all scans from an individual remained within the same Train or Test dataset. The Train dataset comprised of ∼80% of GUSTO and S-PRESTO participants who were not diagnosed with any developmental disorders and assumed to follow normative development. The Test dataset consisted of ∼20% of GUSTO participants with any parent-reported diagnosis in childhood and enriched for early life adversity exposure and high childhood depressive symptoms, 20% randomly stratified S-PRESTO participants, and an independent symptomatic cohort (BRACO-LD). The Train and Test datasets are comparable across age and MRI QC measures (Table S2).

### 2.2 BrainAge model performance

BrainAge models were trained with the the GPBoost algorithm from the gpboost package v1.5.8^30^ predicting chronological age from neuroimaging features with a random effects term for participant ID. Twelve sex-stratified models were trained consisting of different combinations of time-points (2 Model Types: Full-Span vs Phase-Specific models) and neuroimaging features (S1, S2, F1, A1, A2, A3; Fig 1) in the Train dataset. Structural neuroimaging features were 68 cortical thickness and 14 subcortical volume measures. Functional neuroimaging features were a single modularity score representing functional segregation between community clusters, 17 network recruitment measures representing the internal cohesion of a specific network, and 136 network integration measures representing how inter-connected a network was with other networks.

The candidate BrainAge models were then applied to the Test dataset to estimate BrainAge and calculate model metrics for each candidate model. Predicted Age Difference (PAD) was calculated by subtracting the chronological age at scan from predicted BrainAge. Median Absolute Error (MAE) was calculated by averaging the absolute PAD value.

We first examined whether models specific to developmental phase (Phase-Specific: early-and late-childhood) improved in their predictive validity compared to Full-Span models covering the full age range. BrainAge models were assessed in two ways in the Test dataset: (i) model metrics and (ii) the ability of estimated BrainAge to distinguish between healthy and symptomatic subgroups. Overall, the Phase-Specific models had a less skewed median PAD, a lower MAE, and stronger associations with age compared to the Full-Span models (Table 1A-B). The Test dataset was categorized into subgroups based on cohort or cut-offs for different functional measures and subgroup differences were assessed with regression models. The BrainAge predictions from Phase-Specific models showed significant differences for the BRACO learning difficulties cohort (Fig 2A) and high total behavioral problems measured by the Youth Self Report (Fig 2B-C). In addition, the models differed in their ability to distinguish individuals on the basis of cumulative early life adversity (Fig 2D) and high perceived stress (Fig 2E-F). Phase-Specific models were able to distinguish between more subgroups than Full-Span models.

**Figure 2:**
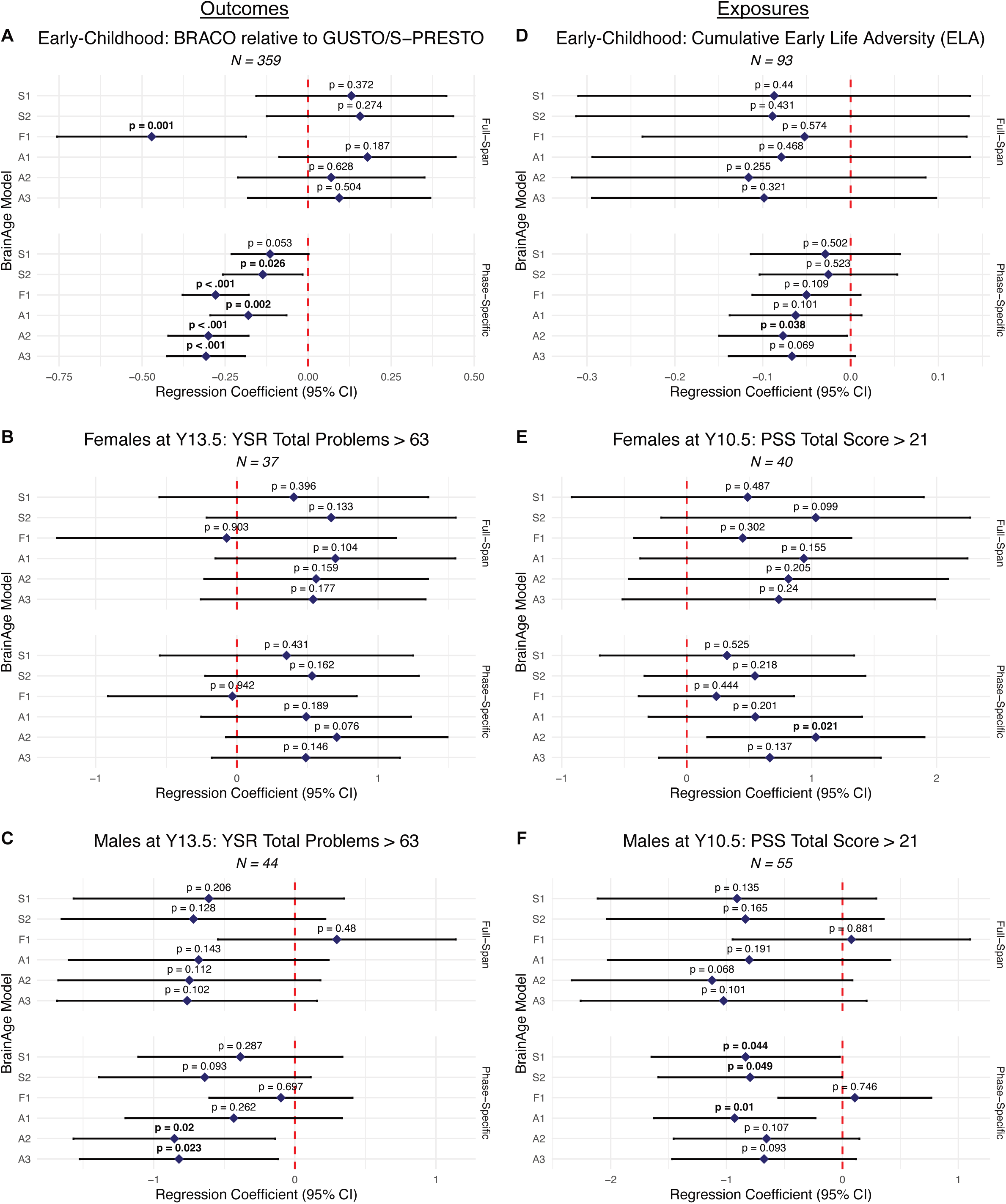
Effect of Predictors on BrainAge estimated from the 12 candidate BrainAge models in the Test dataset. Forest plots show the estimated difference in BrainAge for the following symptomatic subgroups: (A) symptomatic BRACO-LD cohort, high YSR total problems in (B) Females and (C) Males, (D) cumulative ELA, and high perceived stress in (E) Females and (F) Males, relative to the sample norm. Points are the unstandardized regression coefficients adjusted for age, with horizontal error bars indicating 95% Confidence Intervals. The red vertical dashed line at zero indicates the null hypothesis (no difference). Estimates are significant if 95% CI does not cross 0. For example, a negative coefficient (left of the zero line) signifies younger BrainAge in BRACO-LD participants relative to the population reference GUSTO/S-PRESTO. Note: CI, Confidence Interval; PSS, Perceived Stress Scale; YSR, Youth Self Report

**Table 1A:** Summary of model metrics of BrainAge models by type and neuroimaging features for the early-childhood phase.

|  |  | Neuroimaging Features |  |  |  |  |  |
| --- | --- | --- | --- | --- | --- | --- | --- |
| Metric | Type | S1 | S2 | F1 | A1 | A2 | A3 |
| Female |  |  |  |  |  |  |  |
| Median PAD (IQR) | Full-Span | 0.64 (1.96) | 0.57 (2.07) | 1.38 (1.88) | 0.55 (1.94) | 0.69 (1.78) | 0.71 (1.88) |
| Median PAD (IQR) | Phase-Specific | 0.01 (1.59) | -0.06 (1.4) | 0 (1.52) | -0.04 (1.47) | -0.04 (1.27) | -0.04 (1.26) |
| MAE (IQR) | Full-Span | 1.01 (1.39) | 1.02 (1.28) | 1.52 (1.62) | 1.06 (1.25) | 1.02 (1.1) | 0.97 (1.17) |
| MAE (IQR) | Phase-Specific | 0.77 (0.79) | 0.6 (0.79) | 0.77 (0.77) | 0.67 (0.84) | 0.64 (0.72) | 0.63 (0.75) |
| Age r [95% CI] | Full-Span | 0.38 [0.25, 0.5] | 0.33 [0.19, 0.45] | 0.34 [0.2, 0.46] | 0.33 [0.2, 0.45] | 0.38 [0.25, 0.5] | 0.4 [0.27, 0.51] |
| Age r [95% CI] | Phase-Specific | 0.49 [0.38, 0.59] | 0.57 [0.47, 0.66] | 0.41 [0.29, 0.53] | 0.55 [0.44, 0.64] | 0.59 [0.49, 0.68] | 0.59 [0.49, 0.67] |
| Male |  |  |  |  |  |  |  |
| Median PAD (IQR) | Full-Span | 0.75 (1.92) | 0.81 (1.77) | 1.18 (2.13) | 0.9 (1.73) | 0.7 (1.89) | 0.67 (1.81) |
| Median PAD (IQR) | Phase-Specific | 0.02 (1.27) | 0.02 (1.24) | -0.21 (1.6) | 0.07 (1.29) | -0.08 (1.34) | -0.11 (1.24) |
| MAE (IQR) | Full-Span | 1.08 (1.35) | 1.05 (1.23) | 1.36 (1.64) | 1.08 (1.25) | 1.06 (1.35) | 1.06 (1.32) |
| MAE (IQR) | Phase-Specific | 0.66 (0.78) | 0.63 (0.73) | 0.8 (0.87) | 0.64 (0.69) | 0.66 (0.73) | 0.61 (0.75) |
| Age r [95% CI] | Full-Span | 0.41 [0.3, 0.51] | 0.41 [0.3, 0.51] | 0.23 [0.11, 0.34] | 0.41 [0.3, 0.51] | 0.44 [0.33, 0.53] | 0.44 [0.33, 0.53] |
| Age r [95% CI] | Phase-Specific | 0.47 [0.37, 0.56] | 0.52 [0.42, 0.6] | 0.25 [0.13, 0.36] | 0.52 [0.42, 0.6] | 0.48 [0.38, 0.57] | 0.48 [0.38, 0.57] |

Note: statistics shown are median (IQR)

**Table 1B:** Summary of model metrics of BrainAge models by type and neuroimaging features for the late-childhood phase.

|  |  | Neuroimaging Features |  |  |  |  |  |
| --- | --- | --- | --- | --- | --- | --- | --- |
| Metric | Type | S1 | S2 | F1 | A1 | A2 | A3 |
| Female |  |  |  |  |  |  |  |
| Median PAD (IQR) | Full-Span | -0.4 (1.8) | -0.37 (1.85) | -0.89 (3.15) | -0.33 (1.83) | -0.28 (2.06) | -0.33 (2.09) |
| Median PAD (IQR) | Phase-Specific | 0.14 (2.16) | 0.16 (1.9) | -0.12 (3.35) | 0.23 (1.77) | 0.1 (1.97) | 0.09 (1.79) |
| MAE (IQR) | Full-Span | 1.04 (1.22) | 1.02 (1.29) | 1.55 (1.98) | 1.03 (1.25) | 1.09 (1.17) | 1.07 (1.26) |
| MAE (IQR) | Phase-Specific | 1.04 (1.36) | 0.84 (1.28) | 1.63 (1.69) | 0.83 (1.15) | 0.96 (1.05) | 0.87 (1.12) |
| Age r [95% CI] | Full-Span | 0.79 [0.72, 0.85] | 0.82 [0.76, 0.87] | 0.37 [0.22, 0.51] | 0.81 [0.74, 0.86] | 0.8 [0.73, 0.85] | 0.8 [0.73, 0.85] |
| Age r [95% CI] | Phase-Specific | 0.77 [0.68, 0.83] | 0.8 [0.73, 0.85] | 0.38 [0.22, 0.52] | 0.81 [0.74, 0.86] | 0.79 [0.72, 0.85] | 0.81 [0.74, 0.86] |
| Male |  |  |  |  |  |  |  |
| Median PAD (IQR) | Full-Span | -0.44 (2.07) | -0.37 (2.26) | -0.52 (3.01) | -0.42 (2.09) | -0.43 (2.21) | -0.39 (2.11) |
| Median PAD (IQR) | Phase-Specific | 0.03 (2.07) | 0.1 (1.96) | 0.07 (3.37) | 0.03 (1.77) | 0.09 (1.95) | 0.12 (2.01) |
| MAE (IQR) | Full-Span | 1.1 (1.25) | 1.12 (1.31) | 1.37 (2.49) | 1.14 (1.27) | 1.14 (1.28) | 1.16 (1.34) |
| MAE (IQR) | Phase-Specific | 1 (1.03) | 0.98 (1.03) | 1.71 (1.86) | 0.88 (1.17) | 0.99 (1.04) | 1.02 (1.01) |
| Age r [95% CI] | Full-Span | 0.74 [0.66, 0.8] | 0.74 [0.67, 0.81] | 0.42 [0.29, 0.54] | 0.75 [0.67, 0.81] | 0.74 [0.66, 0.8] | 0.74 [0.66, 0.8] |
| Age r [95% CI] | Phase-Specific | 0.75 [0.67, 0.81] | 0.75 [0.68, 0.81] | 0.36 [0.22, 0.49] | 0.76 [0.68, 0.81] | 0.74 [0.66, 0.8] | 0.74 [0.67, 0.81] |

Note: statistics shown are median (IQR)

To account for potential sex-specific effects, we performed sex-stratified regression models to obtain separate estimates in males and females. Estimates for BRACO and cumulative early life adversity showed significant effects in the same direction in males and females (Fig S2). Thus, analyses were combined and estimates for the full cohort were reported. Sex-stratified estimates were reported for perceived stress and YSR total problems. Coefficients estimated for male and female models were in opposite directions (Fig 2B-C, Fig 2E-F). Regression estimates for each subgroup across candidate BrainAge models were reported in Tables S3a-f.

We selected the Phase-Specific “A2” model for subsequent analysis as the model that best distinguished between subgroups enriched for early life adversity, perceived stress, and symptomatic outcomes (Fig 2A, Fig 2C-E). The A2 model consists of cortical thickness, subcortical volumes, functional network integration, and modularity, suggesting that a BrainAge model incorporating both structural and functional features had the best predictive validity.

### 2.3 Neuroimaging Features driving Phase-Specific A2 Model using modularity, functional integration and structural measures

SHapley Additive exPlanations (SHAP) values quantify the importance of each individual feature to the model^31^ where higher values represent greater contribution. We used the percentage of total SHAP contribution (SHAP %) to identify the features that most contributed to each Phase-Specific A2 model stratified by developmental phase and sex (Table S4). Our findings showed that different features drive A2 models specific for the early- and late-childhood phases (Fig 3A-C). For example, the left nucleus accumbens volume is the top contributor for both the female and male early-childhood models (>6.5%) but contributes <0.5% for the late-childhood models (Fig 3B). The thickness of the right lingual gyrus and right medialorbitofrontal cortex are the top contributors for the late-childhood models (Fig 3A). These differences show that different features predict neurodevelopment in the early- and late-childhood phases, underscoring the need for phase-specific BrainAge models.

**Figure 3:**
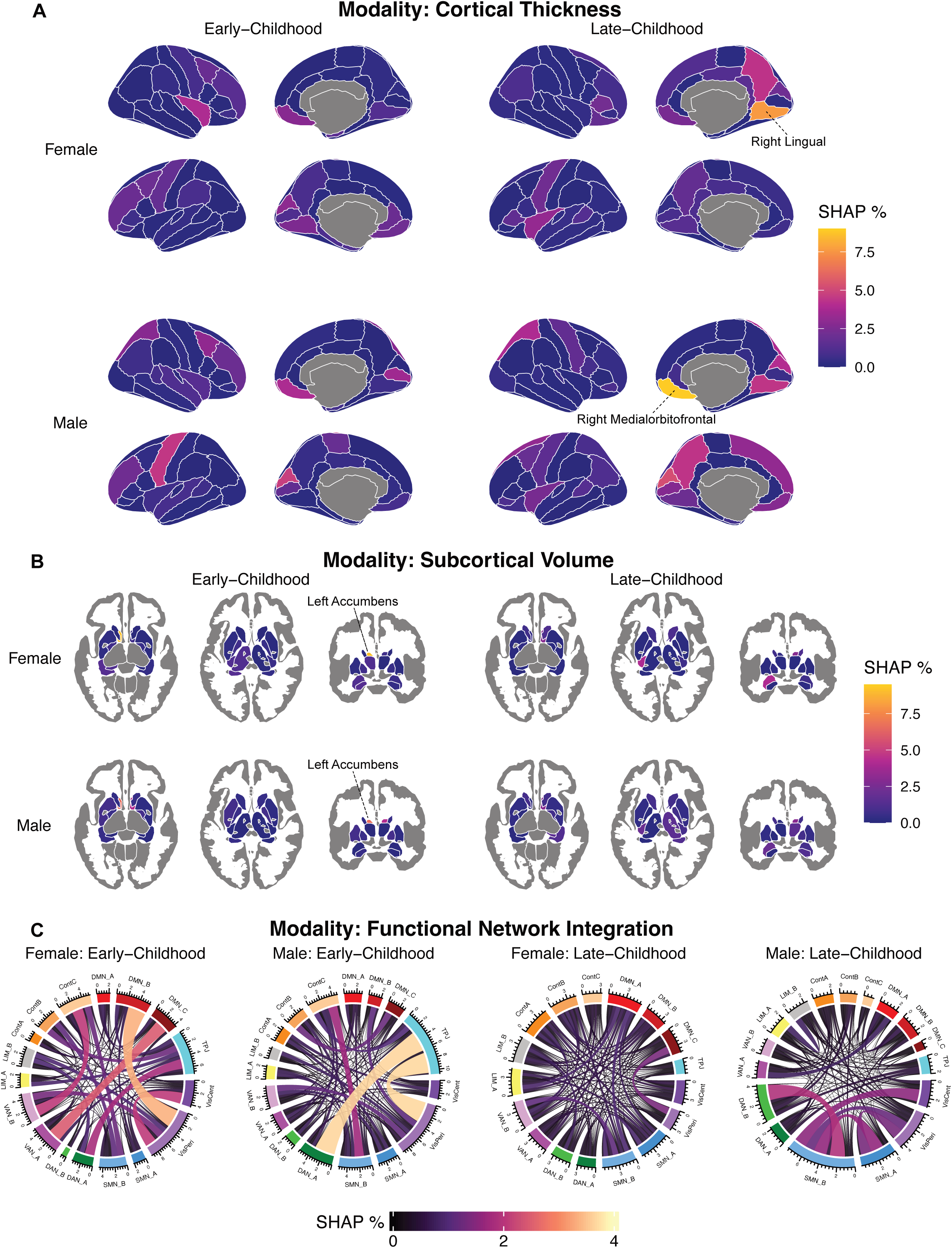
SHAP values for each neuroimaging feature from the selected candidate model (Phase-Specific A2). SHAP values were displayed as a percentage of total SHAP values and split by modality: (A) Cortical thickness, (B) Subcortical volumes, and (C) Functional Network Integration.

### 2.4 BrainAge model benchmarked against other phenotypic age measures

We benchmarked the best performing candidate model (Phase-Specific A2) with four pre-trained BrainAge models from the literature (ENIGMA^7^, CentileBrain^11^, Developmental BrainAge^12^, PyBrainAge^13,32^, Table S5) and two DNA-based biological age measures (PedBE epigenetic clock^33^, telomere length^34^). Biological Age measures were collected at 4 GUSTO time-points and a subset of the BRACO cohort. Biological age measures were only included if DNA was collected within 100 days of the MRI scan. BrainAge estimates were derived for the same Test dataset and assessed using both Median Absolute Error (MAE) and subgroup comparisons. Individuals with Biological Age measures did not overlap fully with the Test dataset and had different sample numbers and thus, were only assessed on subgroup comparisons. Partial Pearson’s correlations between BrainAge and biological age measures controlling for sex are shown in Fig S3.

When benchmarked against published pre-trained models, the Phase-Specific A2 model demonstrated superior predictive precision of age, especially in the early-childhood phase (Fig 4A, Table S6). Specifically, our model achieved an MAE of 0.64 for females and 0.66 for males, compared to the best-performing pre-trained model (Centile: 3.09 Female; 2.68 Male). We observe that pre-trained models show a strongly positive median PAD (median PAD ranging from 2.28 to 15.78 years, Table S6), indicating BrainAge estimates show a right skew that largely over-estimates BrainAge relative to chronological age. This finding is not unexpected given that the training data for these models are largely comprised of adult data.

**Figure 4A:**
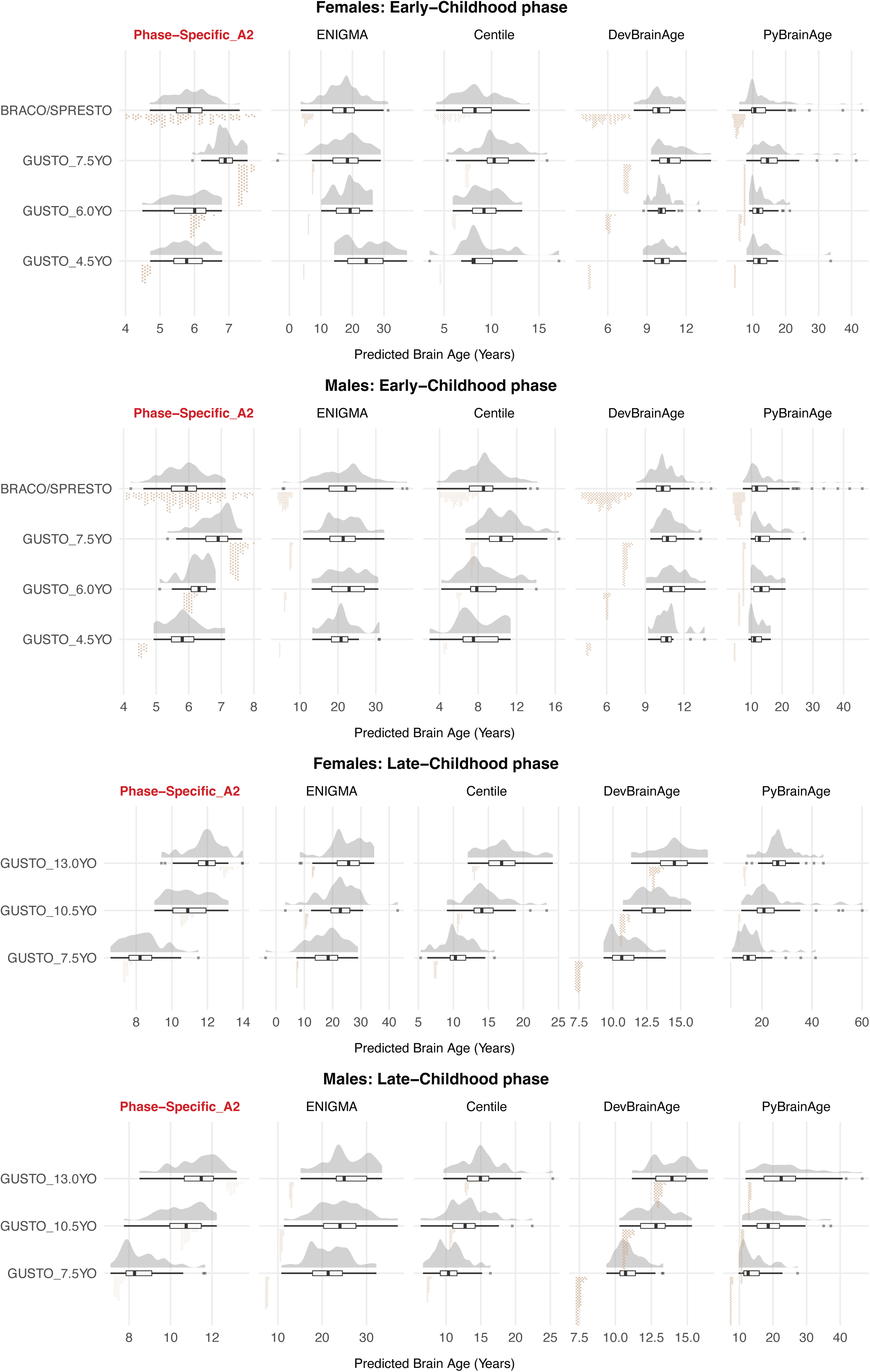
The difference between predicted BrainAge and age at scan of the selected candidate BrainAge model (labelled in red) benchmarked against four published pre-trained BrainAge models. Predicted BrainAge stratified by cohort and time-point is summarised by the horizontal boxplot, with the density plot showing the distribution of the BrainAge estimates for each model. Chronological age at scan is displayed as points (in brown) under the boxplot. The Y axis range is independent to each plot given the differences in the range of age predictions for each model. The distribution of predicted BrainAge from pre-trained models are largely to the left of age at scan, indicating a possible positive bias where the model consistently overestimates age. Plots are faceted by sex and developmental phase (early- and late-childhood).

Furthermore, Phase-Specific A2 exhibited significantly higher clinical sensitivity, capturing significant subgroup differences in more categories than the benchmark models (Fig 4B). We note that when subgroup differences are significant, regression estimates are much larger in pre-trained models, likely reflecting the elevated MAE of the BrainAge estimates and over-estimating the impact of outcomes and exposures on BrainAge. Regression estimates for each subgroup across BrainAge and biological age measures were reported in Tables S7a-f. Our findings support the need for childhood-specific BrainAge models.

**Figure 4B:**
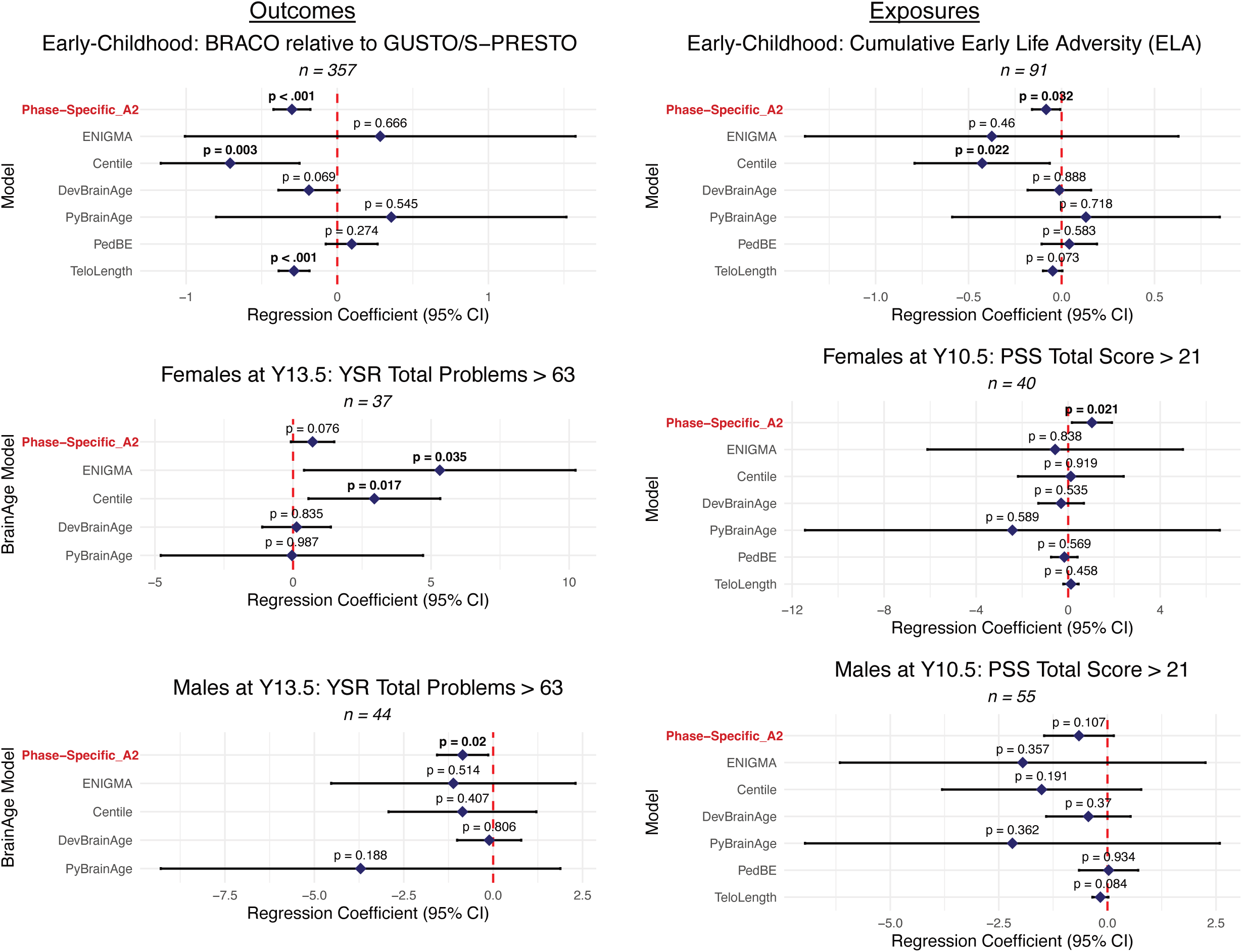
Effect of Predictors on BrainAge estimated from the selected candidate BrainAge model (labelled in red) benchmarked against published pre-trained age proxy measures. Forest plots show the estimated difference in BrainAge for the following symptomatic subgroups: (i) symptomatic BRACO-LD cohort, high YSR total problems in (ii) Females and (iii) Males, (iv) cumulative ELA, and high perceived stress in (v) Females and (vi) Males, relative to the sample norm. Points are the unstandardized regression coefficients adjusted for age, with horizontal error bars indicating 95% Confidence Intervals. The red vertical dashed line at zero indicates the null hypothesis (no difference). Estimates are significant if 95% CI does not cross 0. Note: CI, Confidence Interval; PSS, Perceived Stress Scale; YSR, Youth Self Report

### 2.5 Sensitivity Analyses

We confirmed the robustness of our findings through sensitivity analyses in (i) a non-harmonized Test dataset, (ii) a Test dataset that passed motion criteria, and (iii) Train and Test datasets that passed motion criteria. We observed similar SHAP patterns to the main analysis, with different patterns for the Early-Childhood and Late-Childhood models, when the Phase-Specific A2 model was trained only on scans that passed the motion criteria (Fig S4a). Model metrics were similar across sensitivity analyses (Table S6 & S8), and subgroup regression estimates were similar (Fig S4b).

## 3. Discussion

We showed evidence supporting the need for multi-modal neurodevelopmental BrainAge models aligned to different developmental phases. First, Phase-Specific BrainAge models had lower MAE and were less skewed compared to Full-Span as well as published BrainAge and DNA-based biological age models. Second, neuroimaging features identified as the top contributors for each model were both sex- and phase-specific. Third, a Phase-Specific BrainAge model based on cortical thickness, subcortical volumes, modularity, and functional network integration measures out-performed other candidate models and age proxy measures to distinguish between a range of healthy and symptomatic subgroups. Thus, predicted BrainAge as a biomarker for brain health during development could be improved by developmental phase-specific models that capture both structural and functional changes.

BrainAge models in the developmental context can be a valuable resource as an individual-level predictive biomarker of future outcomes, thus facilitating early identification of neurodevelopmental vulnerability and early intervention. However, there are some additional considerations compared to lifespan/aging models. During childhood, the brain undergoes a series of rapid, non-linear dynamic processes, including synaptic pruning and functional network segregation, that occur within narrow chronological windows^15,17^. We provide evidence for the developmental phase specificity of these processes by showing that different neuroimaging features drive BrainAge models in early- and late-childhood. Our findings are further strengthened by the presence of overlapping individuals across phases, mitigating the confounding effect of individual differences. Thus, pediatric models require higher temporal resolution and models should be trained over smaller age ranges to ensure sensitivity to these developmental transitions. Our findings suggest that the left nucleus accumbens (NAc) volume and functional integration involving the peripheral visual network are key features driving neurodevelopment in early-childhood. Prior literature suggests that reward-related subcortical structures mature earlier than cognitive control-related prefrontal cortical regions^35,36^. The NAc plays an important role in reward processing, shows functional lateralization, and is crucial for executive functioning in children^37^. NAc volume at 6 years of age was also found to mediate the associations between prenatal maternal mental health and attention problems at age 7, suggesting that the NAc at this temporal window represents a scaffold for the developing attention system^38^. Tooley et al. showed that between network connectivity, but not within-network connectivity, showed strong associations with age across childhood, aligning with our finding that our best-performing model (A2) comprised of functional integration measures between networks, but not recruitment within a network^39^. Moreover, they found that positive age effects were most pronounced in the visual network and default mode network, especially in the medial prefrontal cortex^39^, thus highlighting the relevance of these networks for BrainAge models. Our models suggest that cortical thickness, particularly in the frontal lobe and regions associated with visual processing (Table S4), serves as the primary driver of neurodevelopmental maturity in late-childhood. This finding is consistent with longitudinal evidence of synaptic pruning in late-childhood, where the brain undergoes structural fine-tuning to support adult-level function^21,40^. In general, the literature shows that the frontal and occipital regions mature first, while the maturation of association areas that involve information integration from distributed regions extend across adolescence^41,42^. Thus, the features identified in our BrainAge models align with the neurodevelopmental literature.

A second consideration is the interpretation of the BrainAge gap in developmental models. While clinical cohorts typically show a positive Predicted Age Difference (PAD) in aging and adult models that correspond to neurodegeneration or illness-related psychopathology^9^, the interpretation is not so clear-cut in children. In particular, we observed that high perceived stress (Y10.5) and YSR total problems (Y13) had sex-specific effects on estimated BrainAge at that time-point. Specifically, we observed a negative coefficient in males and a positive coefficient in females. In other words, BrainAge estimates were younger in males and older in females in the symptomatic compared to healthy subgroups. These findings were consistent across both functional measures administered at separate time-points. There is a consensus in the literature that sex differences are present in stress-responsivity, partly attributed to the sex-specific exposure to steroid hormones during critical windows of development^43^. Reports of sex differences in stress sensitivity/magnitude and in the brain regions involved provide an explanation for our sex-specific findings^44,45^. Another possibility is the effect of puberty, which is of particular relevance in late-childhood and shows sex differences in both timing and tempo with implications for emergent behavioral problems^46^. Thus, it is difficult to judge whether a positive or negative PAD is beneficial or harmful in developmental models. The PAD reflects the brain’s adaptive response to different environmental demands^47^ and findings should be interpreted within a suitable context as well as with respect to specific functional outcomes. This complication of developmental models was also noted by Whitmore & Beck^48^.

Our findings suggest that neuroimaging-based BrainAge estimates captured subgroup differences better than DNA-based biological age measures, at least for the range of exposures/outcomes we assessed. We note modest associations between Phase-Specific A2 predicted BrainAge and the PedBE (r = 0.27, p = 0.016) and telomere length (r = −0.23, p = 0.033) in the BRACO-LD cohort (Fig S3). While the associations between BrainAge and DNA-based biological age were outside the scope of the current study, it would be interesting to explore which outcomes are best captured by different developmental biomarkers in future studies.

Study limitations must be considered. First, BrainAge gap estimates reflect model prediction errors and can partially be attributed to noise, such as poor-quality scans due to head motion, instead of meaningful biological variance. We performed sensitivity analyses for motion and data harmonization to assess the robustness of our findings and study conclusions remain unchanged. Second, the sample size is modest for subgroup comparisons within the Test dataset. We accounted for this by benchmarking our BrainAge estimates against both pre-trained BrainAge models from the literature and DNA-based biological age measures. The question then becomes whether our Phase-Specific BrainAge models are comparable to published models. Third, our longitudinal cohorts were designed to collect data in waves at different time-points. Hence, scan data were clustered around narrow age ranges and not spread across the whole childhood. Our models were also trained on Singaporean pediatric cohorts, and it is possible that study findings (for example, the key neuroimaging features identified at each phase) may not be generalizable to other cohorts of different genetic ancestry. We acknowledge these limitations and position this paper as a proof-of-concept study to inform the development of childhood neurodevelopmental models. Future iterations should involve integrating diverse, publicly available international neuroimaging datasets to ensure broader generalizability. A larger sample size will allow for an expanded analysis of longitudinal associations to study the rate of change of BrainAge measures beyond cross-sectional subgroup comparisons.

In conclusion, we present a proof-of-concept for developmental models supporting BrainAge models of higher temporal resolution that align to different childhood developmental phases. Our candidate BrainAge model provided superior sensitivity to the subtle neuroanatomical variations associated with a range of outcomes, including learning difficulties, stress and behavioral problems. By leveraging internal training data that closely mirrors the developmental characteristics of our cohort, our BrainAge models achieved a higher degree of predictive precision and clinical utility.

## 4. Methods

### 4.1 Cohorts

Our data were drawn from the participants of three independent, Singaporean pediatric cohorts. The Growing Up in Singapore Towards healthy Outcomes (GUSTO) study^27,28^ (recruitment: June 2009 to September 2010) and the Singapore Preconception Study of Long-Term Maternal and Child Outcomes (SPRESTO) study^29^ (recruitment: February 2015 to October 2017) are population-based birth cohorts involving multiple waves of data collection from mother-child dyads. Data collection time-points range from pregnancy to adolescence and include environmental exposures, behavioral measures and biological samples. In GUSTO, longitudinal neuroimaging data was collected at 5 time-points (Y4.5, Y6.0, Y7.5, Y10.5, and Y13.0 years of age; n = 671, 1792 scans). In S-PRESTO, neuroimaging data was collected once between ages 4 to 6 years (n = 120). The BRAin COnnectivity and Learning Difficulties (BRACO-LD) study is an ongoing pediatric symptomatic cohort (recruitment: 2023 to 2026). Children with learning difficulties were recruited through clinic referrals and from community sources. Neuroimaging data was collected once between ages 4 to 8 years (n = 214). The S-PRESTO and BRACO-LD time-points were designed to have cross-cohort compatibility and pre-harmonized prior to data collection. All three studies were approved by the relevant Singaporean Institutional Review Boards (Supplement 1) and all study investigations accord with the principles expressed in the Declaration of Helsinki. Written informed consent was obtained from all guardians on behalf of the children enrolled in this study. This study followed the Strengthening the Reporting of Observational Studies in Epidemiology (STROBE) reporting guidelines for cohort studies^49^. Fig 1 depicts the data collection time-points. Demographics and MRI quality check (QC) measures across the cohorts and time-points are summarized in Tables S1a-b.

### 4.2 Dataset Split

The total sample (n = 1005, 2126 scans) was split into a Train (n = 633, 1504 scans) and Test dataset (n = 372, 622 scans). The Train dataset comprised of ∼80% of GUSTO and S-PRESTO participants who were not diagnosed with any developmental disorders and assumed to follow normative development. The Test dataset consisted of ∼20% of GUSTO participants with any parent-reported diagnosis in childhood and enriched for early life adversity exposure and high childhood depressive symptoms, 20% randomly stratified S-PRESTO participants, and an independent symptomatic cohort (BRACO-LD). The Train and Test datasets are compared in Table S2.

### 4.3 MRI Acquisition and Processing

Supplement 1 provides detailed information on acquisition parameters and processing steps. T1-weighted and resting state functional MRI (rs-fMRI) scans were collected across three sites: GUSTO Y4.5 and Y6.0 (site 1, Siemens 3-Tesla Magnetom Skyra); GUSTO Y7.5, Y10.5 and Y13 (site 2, Siemens 3-Tesla Magnetom Prisma) and SPRESTO and BRACO-LD (site 3, Siemens 3-Tesla Prisma Fit).

T1-weighted scans were processed with the default FreeSurfer v7.1.1 recon-all pipeline^50^. We extracted 68 bi-lateral cortical thickness measures from the Desikan-Killiany cortical parcellation and 14 subcortical volume measures from seven bi-lateral structures from the aseg parcellation. Volume measures were scaled to estimated total intracranial volume. The Euler number was calculated as a quality control measure for FreeSurfer output.

rs-fMRI data was processed as previously described, with the default pre-processing and de-noising pipelines using the CONN Toolbox v20b^51^. Scan volumes with framewise displacement above 0.9mm or global BOLD signal changes above 5 standard deviations were identified as potential outliers. Functional data was smoothed using spatial convolution with a Gaussian kernel of 6mm full width half maximum (FWHM), and the first four scans excluded to allow for magnetic field saturation.

Regions of Interest (ROIs) were 400 cortical regions^52^ assigned to the 17 functional networks identified by Yeo et al^53^. For each scan, 400 x 400 functional connectivity (FC) matrices were computed by measuring the bivariate correlation coefficients of the BOLD time series between each seed and target ROIs through a haemodynamic response factor (hrf)-weighted general linear model. Brain topology measures were computed as previously described^54–56^. A tuned Louvain community detection algorithm was applied to FC matrices generating a community assignment for each ROI^57^. This analysis was averaged across 100 iterations for stability. A modularity score was calculated that represents functional segregation between community clusters. An allegiance matrix was then generated for each of the 400 ROIs to one of the 17 functional networks. Network integration is defined as the probability of a region being assigned to the same community as regions from another network. Network recruitment is defined as the probability of a region being assigned to the same community as other regions from the same network. Measures per region were averaged at the network level to obtain 136 between-network integration measures and 17 network recruitment measures. Network topology measures were normalised using the distribution from 10,000 iterations with randomly permuted functional connectivity matrices.

As GUSTO data was collected at two sites (Site 1: Y4.5 and Y6.0; Site 2: Y7.5, Y10.5, Y13) and S-PRESTO and BRACO-LD data were collected at a 3^rd^ site, all neuroimaging measures were harmonized across site with longitudinal ComBat v0.0.0.90^58^. A sensitivity analysis was performed with scans that passed motion criteria (<25% outlier volumes, Euler value < 25^th^ percentile – 1.5*IQR).

### 4.4 Environmental/Behavioral Measures collected in GUSTO

*Cumulative ELA:* We calculated a cumulative adversity score that encapsulated early life adversity (ELA) conditions, each of which is known to increase the risk for mental health disorders in the epidemiological literature^59–61^. Participants scored 1 point for each early life adversity factor (e.g., maternal health, socio-economic variables) for which they met criteria^62^ (Table S9). A higher score indicated greater cumulative ELA.

*PSS:* The Perceived Stress Scale-10 (PSS)^63^, administered at age 10 years, is a self-reported questionnaire of ten items measuring perceptions of recent stress where higher scores indicate greater perceived stress. Individuals scoring above the 90^th^ percentile (PSS total score > 21) were categorized as the high stress subgroup.

*YSR:* The Youth Self Report (YSR) was administered at age 13 years. The YSR is a self-reported questionnaire of 112 items assessing social, emotional, and behavioral problems^64^. High YSR total problems is a predictor of poor mental health outcomes^65^. Individuals with YSR total problems T-score > 63^66^ were categorized as the high YSR subgroup.

### 4.5 Age Proxy Measures

BrainAge measures from four pre-trained models from the literature were derived: ENIGMA-Brain Age^7^, CentileBrain^11^, Developmental Brain Age^12^, PyBrainAge^13,32^. Input features for all four models were FreeSurfer extracted morphometric measures following developers’ protocol (Supplement 1). Models largely covered the lifespan, apart from the Developmental Brain Age model which was trained on youths aged 9-19 years (Table S5).

Two Biological Age measures were calculated from DNA extracted from buccal swabs. The Pediatric-Buccal-Epigenetic (PedBE) epigenetic clock is largely specific to pediatric buccal epithelial cells and is derived from 94 CpG methylation sites^33^. Telomere length was determined using a validated quantitative polymerase chain reaction (qPCR) protocol measuring the relative ratio of telomere and β-globin gene copy numbers^34^. Telomere length shortens with each cell division, with shorter telomeres associated with increased biological age.

### 4.6 Statistical Analysis

All statistical analyses were performed in R (v4.3.0)^67^. Alpha level was set at p<0.05 (two-tailed) unless otherwise specified.

*Candidate BrainAge models:* 12 sex-stratified models were performed consisting of different combinations of time-points (2 Model Types: Full-Span vs Phase-Specific models) and neuroimaging features (S1, S2, F1, A1, A2, A3; Fig 1).

The GPBoost algorithm from the gpboost package v1.5.8^30^ was used to train BrainAge models predicting chronological age from neuroimaging features with a random effects term for participant ID. GPBoost combines tree-boosting and Gaussian processes with mixed-effects models, allowing for the predictive power of machine learning models in correlated longitudinal data^68^.

We employed 5-fold repeated cross-validation with 5 repeats to optimize model hyperparameters within the Train dataset. Partitioning was grouped by participant ID to ensure that all scans from an individual remained within the same internal training or validation set. In each iteration, 20% of the Train participants were held out for validation, and optimal hyperparameters were selected by minimizing Root Mean Square Error (RMSE) with a grid search approach. The final model was then retrained on the full Train dataset and applied to the Test dataset to predict BrainAge.

Model metrics were calculated for the Test dataset. Predicted Age Difference (PAD) was calculated by subtracting the chronological age at scan from the predicted age. Median Absolute Error (MAE) was calculated by averaging the absolute PAD value.

Shapley Additive exPlanations (SHAP) values were obtained with the SHAPforxgboost package v0.1.3^69^. SHAP values represent a feature’s contribution to the change in the model output relative to the average baseline prediction^31^ where higher values represent greater contribution. Feature importance was expressed as a percentage of total SHAP contribution, where the absolute SHAP value of each feature was divided by the sum of all feature contributions.

*Model Performance:* We assessed model performance by integrating predictive accuracy and bias (MAE, PAD skewness) with an evaluation of clinical utility across symptomatic subgroups. We first compared internal candidate models. We then selected the best-performing BrainAge model and benchmarked it against pre-trained BrainAge models from the literature and biological age measures. For comparative consistency, BrainAge estimates were evaluated within the same Test dataset subsample, stratified by developmental phase and sex.

To assess whether BrainAge measures reflected environmental and functional differences, linear regression models were performed according to the following equation:

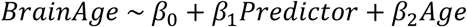

where Predictor represents subgroup membership for (i) symptomatic BRACO-LD cohort, (ii) cumulative ELA, (iii) high stress, and (iv) high YSR total problems. Subgroups were binary variables, except for cumulative ELA which is an ordinal variable (0 to 5). Subgroup details are in Supplement 1. Age was included as a nuisance covariate to ensure that observed differences reflected BrainAge differences rather than variance in actual age. BrainAge estimates closest to the predictor time-points were used.

To account for potential sex-specific effects, we performed sex-stratified regression models to obtain separate estimates in males and females. Male and female datasets were combined if significant estimates were not in opposite directions to increase statistical power. Sex-stratified regression estimates were reported if significant estimates were in opposite directions in males and females.

Given the focus of the analysis on comparative benchmarking and not hypothesis-testing and the recognized limitations of statistical power in smaller neuroimaging cohorts, nominal p-values were reported to avoid an undue inflation of Type II error rates. Instead, we adopted an estimation-based approach, reporting 95% Confidence Intervals alongside effect sizes.

## Data Availability

The data that support the findings of this study are not publicly available. Restrictions apply to the availability of these data, which were used under license for the current study. Data is, however, available from the corresponding author upon reasonable request and with the permission of the A*STAR Institute for Human Development and Potential.

## Code Availability

Study analyses were carried out in R using publicly available R packages. R code (model training, validation, and visualization of findings) is publicly available in the BrainAge-Childhood repository on Github at https://github.com/sychan-code/BrainAge-Childhood. The code is distributed under the Apache License 2.0.

## Author Contributions

Conceptualization was done by SYC. Data processing was carried out by SYC, PH, ALT, AN, JL, ZMN, JSMC, AMAM, and XYHL. Formal analysis and manuscript preparation was performed by SYC. Review and editing were carried out by PH, ALT, AN, JSMC, HJZ, BTTY, YSC, PG, JE, RD, DW, MJM, and APT. Supervision was provided by MJM and APT. Project administration and funding acquisition were provided by MVF, YSC, PG, JE, MJM, and APT.

## Acknowledgements

The study is supported by the National Research Foundation (NRF) under the Open Fund-Large Collaborative Grant (OF-LCG; MOH-000504) administered by the Singapore Ministry of Health’s National Medical Research Council (NMRC) and the Agency for Science, Technology and Research (A*STAR). In RIE2025, GUSTO and S-PRESTO are supported by funding from the NRF’s Human Health and Potential (HHP) Domain, under the Human Potential Programme. BRACO-LD is supported by funding from the NMRC Transition Award (MOH-001273-00).

SYC is supported by funding from the NMRC Open Fund – Young Individual Research Grant (MOH-001149-00). PH is supported by funding from the NMRC Open Fund – Young Individual Research Grant (MOH-001857-00). BTTY is supported by the NUS Yong Loo Lin School of Medicine (NUHSRO/2020/124/TMR/LOA), the Singapore National Medical Research Council (NMRC) LCG (OFLCG19May-0035), NMRC CTG-IIT (CTGIIT23jan-0001), NMRC OF-IRG (OFIRG24jan-0006; OFIRG24jul-0049), NMRC STaR (STaR20nov-0003), Singapore Ministry of Health (MOH) Centre Grant (CG21APR1009), the United States National Institutes of Health (R01MH133334 & 2R01MH120080) and the Singapore National Research Foundation (NRF) Investigatorship (NRFI10-2024-0014). MJM is supported by funding from the Hope for Depression Research Foundation, USA, the Jacobs Foundation, Switzerland, the NRF and the Agency for Science Technology and Research (A*STAR), Singapore, under its Prenatal/Early Childhood Grant (Grant No. H22P0M0001). APT is supported by funding from the NMRC Transition Award (MOH-001273-00). Any opinions, findings and conclusions or recommendations expressed in this material are those of the authors and do not reflect the views of the funders.

