## Supplement 1; Fig S1; Fig S2; Fig S3; Fig S4a-b; Tables S1a-b; Tables S2; Tables S3a-f; Table S4; Table S5; Table S6; Tables S7a-f; Table S8; Table S9 for "Developing a multi-modal neuroimaging-based BrainAge model across childhood"

Supplement 1: Supplementary Methods

Supplementary Fig S1: Study flowchart and analysis numbers

Supplementary Fig S2: Sex-stratified regression estimates for BRACO and cumulative early life adversity

Supplementary Fig S3: Partial Pearson's correlations between BrainAge and biological age measures

Supplementary Fig S4a-b: Sensitivity Analyses for MRI Motion and Harmonization

Supplementary Tables S1a-b: Summary of demographic and MRI QC measures by cohort and time-points

Supplementary Table S2: Age and MRI QC measures stratified by Train/Test and time-point

Supplementary Tables S3a-f: Regression estimates across candidate BrainAge models

Supplementary Table S4: Top contributing regions based on SHAP values for selected model Phase-Specific A2

Supplementary Table S5: Details of 4 pre-trained BrainAge models from the literature

Supplementary Table S6: Model performance metrics across selected A2 BrainAge model with pre-trained BrainAge models (n = 731)

Supplementary Table S7a-f: Regression estimates for selected A2 BrainAge model and published pre-trained BrainAge and biological age measures

Supplementary Tables S8: Model performance metrics across selected A2 BrainAge model with pre-trained BrainAge models for motion subset (n = 638)

Supplementary Table S9: Adversity Score Calculation

Supplement 1: Supplementary Methods

### Cohorts

Study flow chart and analysis numbers are shown in Supplementary Fig S1.

GUSTO

The GUSTO study recruited pregnant women aged 18 years and above, attending their first trimester antenatal dating ultrasound scan clinic at Singapore's two major public maternity units, National University Hospital (NUH) and KK Women's and Children's Hospital (KKH), between June 2009 and September 2010. Participants were (i) Singapore citizens or permanent residents who were of (ii) Chinese, Malay or Indian ethnicity with homogeneous parental ethnic background, who (iii) had the intention of eventually delivering in NUH or KKH and (iv) intended to reside in Singapore for the next 5 years. Furthermore, (v) only women who agreed to donate birth tissues (including cord, placenta and cord blood) at delivery were included. Mothers receiving chemotherapy, psychotropic drugs or who had type I diabetes mellitus were excluded.

The GUSTO study was approved by the National Healthcare Group Domain Specific Review Board (D/2009/021 and B/2014/00411) and the SingHealth Centralized Institutional Review Board (D/2018/2767 and A/2019/2406).

The GUSTO study is registered under the ClinicalTrials.gov ID NCT01174875.

Out of the 671 GUSTO participants included in the study, 35 individuals had a parent-reported diagnosis of one or more of the following: (1) ASD, (2) ADHD/Attention Problems, (3) Behavioural/Conduct/Anger Management Problems, (4) Developmental Delays, (5) Emotional Difficulties, (6) Anxiety/Selective Mutism, (7) Dyslexia, (8) Neurological/spinal issues, (9) Diagnosed epilepsy or history of repeated non-fever related seizures or on anti-epileptic medication, (10) Unspecified behavioural/attention issues but seeing psychiatrist/psychologist. These individuals were assigned to the Test dataset.

S-PRESTO

The S-PRESTO study recruited women, aged 18 to 45 years, between February 2015 and October 2017. An upper age limit of 35 years was introduced from mid-July 2016 onward to increase the likelihood that recruited women conceive within the time frame of the study. Other inclusion criteria include: participants were (i) planning to conceive within 1 year of recruitment, (ii) intending to reside in Singapore for the next 5 years, (iii) of Chinese, Malay, Indian ethnicity or any combinations of these 3 ethnic groups, and (iv) able to provide written, informed consent. Women who did not become pregnant after > 12 months from recruitment, had pregnancy complications, had been actively trying to conceive for > 18 months, were pregnant at recruitment, with pre-existing diabetes, or on certain medications in the past month (systemic steroids, anticonvulsants, HIV/Hepatitis B/Hepatitis C medication) were excluded.

The SPRESTO study was approved by the National Healthcare Group Domain Specific Review Board and the SingHealth Centralized Institutional Review Board (reference 2014/692/D).

The S-PRESTO study is registered under the ClinicalTrials.gov ID NCT03531658.

BRACO-LD

The BRACO-LD study recruited children aged 4 to 8 years with functional learning difficulties between November 2021 and June 2026. Recruitment was done through referral from child development units at the Singapore National University Hospital (NUH) and community outreach such as social media advertisements. Children were included if they had a known diagnosis of learning difficulties or were participating in formal interventions. Children without a known diagnosis or attending interventions were screened using a comprehensive test battery and included only if they scored below -1 SD in any cognitive domain. Exclusion criteria included: (1) inability to tolerate MRI scanning or psychometric evaluation, (2) history of medical conditions affecting learning ability, and (3) presence of metallic implants.

An initial pilot study was approved by the National Healthcare Group Domain Specific Review Board (DSRB- 2021-00514). The main study was subsequently approved by the National Healthcare Group Domain Specific Review Board (DSRB-2022-00792), the National University of Singapore Institutional Review Board (NUS-IRB-2022-185), and the A*STAR Institutional Review Board (ASTAR-IRB-2024-108).

### MRI Acquisition Parameters

During the acquisition of neuroimaging data, children were instructed to lie still without watching TV. All children were scanned in a tertiary pediatric facility with staff experienced in handling young children. Children also underwent training in a mock scanner.

<u>Site 1 (Magnetom Skyra, Siemens)</u>
3D T1-weighted Magnetized Prepared Rapid Gradient Echo (MPRAGE) images were acquired with the following imaging parameters: repetition time = 2000ms, echo time = 2.08ms, inversion time = 877ms, flip angle = 9, field of view = 192mm, matrix size = 192 x 192, a total of 160 contiguous sagittal slices were acquired with 1mm isotropic voxels, slice thickness = 1mm.

Resting state fMRI: Resting-state functional MRI (rsfMRI) images were acquired with a gradient-echo planar imaging sequence sensitive to blood oxygenation level-dependent (BOLD) contrast. One run of rsfMRI was collected: 5.32 minutes (120 volumes), repetition time = 2660ms, echo time = 27ms, flip angle = 90 degrees, 3mm isotropic voxels, matrix = 64 x 64, field of view = 192mm, 48 interleaved axial slices with 3mm slice thickness and no gap.

<u>Site 2 (Magnetom Prisma, Siemens)</u>
3D T1-weighted Magnetized Prepared Rapid Gradient Echo (MPRAGE) images were acquired with the following imaging parameters: repetition time = 2000ms, echo time = 2.08ms, inversion time = 877ms, flip angle = 9, field of view = 192mm, matrix size = 192 x 192, a total of 192 contiguous sagittal slices were acquired with 1mm isotropic voxels, slice thickness = 1mm.

Resting state fMRI: Resting-state functional MRI (rsfMRI) images were acquired with a gradient-echo planar imaging sequence sensitive to blood oxygenation level-dependent (BOLD) contrast. One run of rsfMRI was collected: Repetition time = 2620ms, echo time = 27ms, flip angle = 90 degrees, 3mm isotropic voxels, matrix = 64 x 64, field of view = 192mm, 48 interleaved axial slices with 3mm slice thickness and no gap. Depending on the time-point, the number of scan volumes collected was: 120 volumes (Y7.5), 183 volumes (Y10.5), 114 volumes (Y13).

<u>Site 3 (Magnetom Prisma Fit, Siemens)</u>
3D T1-weighted Magnetized Prepared Rapid Gradient Echo (MPRAGE) images were acquired with the following imaging parameters: repetition time = 2000ms, echo time = 2.08ms, inversion time = 877ms, flip angle = 9, field of view = 192mm, matrix size = 192 x

192, a total of 192 contiguous sagittal slices were acquired with 1mm isotropic voxels, slice thickness = 1mm.

Resting state fMRI: Resting-state functional MRI (rsfMRI) images were acquired with a multi-band gradient-echo planar imaging sequence sensitive to blood oxygenation level-dependent (BOLD) contrast. One run of rsfMRI was collected: 8.00 minutes (668 volumes), repetition time = 719ms, echo time = 30ms, flip angle = 50 degrees, 2.5mm isotropic voxels, matrix = 88 x 88, field of view = 220mm, 60 interleaved axial slices with 2.5mm slice thickness and no gap with a multi-band acceleration factor of 6.

### MRI preprocessing

Raw DICOM format images were converted into NIFTI format using the dcm2niix tool[1].

T1-weighted MPRAGE: Pre-processing was performed using FreeSurfer v7.1.1 with the default recon-all pipeline (http://surfer.nmr.mgh.harvard.edu).
The automated pipeline included motion correction, skull stripping, B1 bias field correction, and automated transformation into a conformed space. Cortical surfaces were reconstructed to delineate the gray-white matter boundary (white surface) and the pial surface. Cortical thickness was calculated as the shortest distance between these two surfaces at each vertex across the mantle.

Resting state fMRI: Pre-processing was performed using the CONN toolbox v20b with the default pre-processing pipeline [2]. Briefly, scans underwent realignment with SPM12 realign & unwarp procedure [3], where all scans were coregistered and resampled to the first scan using b-spline interpolation. Temporal misalignment between different slices of the functional data was corrected using SPM12 slice-timing correction (STC) procedure [4], where the functional data is time-shifted and resampled using sinc-interpolation to match the time in the middle of each acquisition time. Scan volumes with framewise displacement above 0.9mm or global BOLD signal changes above 5 standard deviations were identified as potential outliers. Framewise displacement was computed at each timepoint by estimating the largest displacement among six control points placed at the center of a defined bounding box. Global BOLD signal change was calculated as the change in average BOLD signal within SPM's global-mean mask scaled to standard deviation units at each timepoint. Functional and anatomical data were normalized into standard MNI space and segmented into grey matter, white matter, and CSF tissue classes using SPM12 unified segmentation and normalization procedure [5]. Both functional and anatomical data were resampled using 4th order spline interpolation. Functional data was smoothed using spatial convolution with a Gaussian kernel of 6mm full width half maximum (FWHM), and the first four scans excluded to allow for magnetic field saturation. For denoising, BOLD signal variance over time explained by nuisance variables was removed from the data using Ordinary Least Squares regression. Noise components include identified outliers, motion parameters, and mean white matter and CSF signal [6]. Next, BOLD time series were band-pass filtered to preserve only frequencies between 0.008 and 0.09 Hz [7,8].

### MRI Measures

Cortical Thickness and Subcortical Volumes:
Cortical Thickness: 68 ROIs from the Desikan-Killiany cortical parcellation (both left and right hemisphere).

Subcortical Volume: 14 ROIs from 7 bi-hemisphere subcortical regions from the default FreeSurfer aseg parcellation – Thalamus, Caudate, Putamen, Pallidum, Hippocampus,

Amygdala, Nucleus Accumbens. Volumes for each ROI were scaled to represent the % volume of estimated total intracranial volume.

Functional Connectivity:
Regions of Interest (ROIs) were 400 cortical regions assigned to the 17 functional networks identified by Yeo et al (Visual Central, Visual Peripheral, Somatomotor A, Somatomotor B, Dorsal Attention A, Dorsal Attention B, Salience/Ventral Attention A, Salience/Ventral Attention B, Limbic A, Limbic B, Fronto-Parietal/Control A, Fronto-Parietal/Control B, Fronto-Parietal/Control C, Default Mode Network A, Default Mode Network B, Default Mode Network C, Temporal Parietal)[9,10].

For each scan, functional connectivity (FC) matrices were computed by measuring the bivariate correlation coefficients of the BOLD time series between each seed and target ROIs through a haemodynamic response factor (hrf)-weighted general linear model.

FC matrices were then used to compute brain network topology measures. A tuned Louvain community detection algorithm[11] was used to cluster the ROIs into community clusters by generating a data-driven group assignment for each ROI. This analysis was averaged across 100 iterations for stability. A modularity score was calculated that represents the degree of segregation between the different Louvain community clusters.

Subsequently, the group assignments were used to create an allegiance matrix for each of the 400 ROIs to one of the 17 functional networks, generating integration and recruitment coefficients. The integration coefficient for each region is defined as the probability of a region being assigned to the same community as regions from another network. The recruitment coefficient for each region is defined as the probability of a region being assigned to the same community as other regions from the same network. Integration and recruitment measures were averaged at the network level to obtain 136 between-network integration measures and 17 network recruitment measures. Network topology measures were normalised using the distribution from 10,000 iterations with randomly permuted functional connectivity matrices to account for differences in the number of regions for each network.

Harmonization:
As GUSTO data was collected a two sites (Site 1: Y4.5 and Y6.0; Site 2: Y7.5, Y10.5, Y13) and S-PRESTO and BRACO-LD data were collected at a 3$^{rd}$ site, all neuroimaging measures were harmonized across site with longitudinal ComBat v0.0.0.90[12].

### MRI motion

To maximize power in our analysis, subjects were not excluded from further analysis based on motion parameters. Instead, a sensitivity analysis was conducted in participants who passed motion criteria for both structural and functional connectivity data to determine if motion changed the main findings.

Structural Data
For cortical thickness and subcortical volumes, the Euler number was calculated to assess cortical surface reconstruction quality and scan data quality. The Euler number is derived from a FreeSurfer output (x, the number of surface holes) using the formula:
$2 – 2x$.

A value closer to 2 indicates better quality, while more negative values indicate topological defects (holes) often caused by noise or motion artifacts. Scans with euler values < 25$^{th}$ percentile – 1.5*IQR (per time-point) are flagged, and excluded if they failed a visual QC of the FreeSurfer outputs. 43 scans failed the structural data criteria and were excluded in the sensitivity analyses.

Functional Connectivity
For functional measures, we extracted the three motion QC measures generated by the CONN toolbox pipeline – mean motion (QC_MeanMotion - mean relative motion over all volumes), max motion (QC_MaxMotion - max relative motion over all volumes), invalid scans (QC_InvalidScans - scan volumes that exceeded the motion threshold and are labelled as outliers).

Scans that exceeded a threshold of >25% invalid scans were excluded. 289 scans failed the functional data criterion and were excluded in the sensitivity analyses.

### Sensitivity Analyses

Sensitivity analyse were conducted to confirm the robustness of the main findings for the selected Phase-Specific A2 model:

1. Non-harmonized Test Dataset – Phase-Specific A2 applied to raw neuroimaging features before longitudinal ComBat.
2. Motion subset (Test Dataset) – Model performance reported only for subset of Test Dataset that passed motion criteria
3. Motion subset (Train + Test Dataset) – Phase-Specific A2 model (M) was generated only on the Train Dataset scans that passed motion criteria. This Phase-Specific A2 model (M)-version model was then applied to the motion subset Test Dataset. Model performance was assessed.

Motion Subset Sample Size
Total: n = 934 IDs, 1822 scans

Train:
Females (Early): n = 255, 396 scans
Females (Late): n = 229, 467 scans
Males (Early): n = 234, 336 scans
Males (Late): n = 223, 404 train

Test:
Females (Early): n = 131, 158 scans
Females (Late): n = 54, 117 scans
Males (Early): n = 192, 222 scans
Males (Late): n = 70, 141 scans

### Established Brain Age models from the literature

Input features for the models were FreeSurfer extracted morphometric measures (cortical thickness, cortical surface area, subcortical volumes) from the default FreeSurfer parcellations (Desikan- Killiany, Destrieux, aseg). Input features were prepared according to the feature list and naming convention specified by the model developer. No additional data scaling or harmonization steps were applied for these models.

ENIGMA BrainAge model[13]: Scripts provided by the ENIGMA consortium (https://github.com/ENIGMA-git/ENIGMA-FreeSurfer-protocol?tab=readme-ov-file) were used for data preparation to ensure column names aligned with the input features needed for the models. Sex-stratified input features were uploaded to the PHOTON platform (https://photon-ai.com/enigma_brainage) for calculation of ENIGMA BrainAge outputs.

CentileBrain model[14]: Sex-stratified input features were uploaded to the Centile platform (https://centilebrain.org/#/brainAGE2) for calculation of CentileBrain outputs. The predicted age output was used.

Developmental Brain Age[15]: Model was applied according to the scripts and model provided in the github page (https://github.com/GitDro/DevelopmentalBrainAge).

PyBrainAge[16,17]: PyBrainAge was calculated using the helper functions and the model provided from the PyBrainAge OSF page (https://osf.io/mxueh/wiki?wiki=734gh)

**DNA-based biological age**

Buccal swabs were collected, and a sub-sample that was collected within 100 days of the MRI scan was sent for DNA extraction and further analysis for the GUSTO cohort (Y4.5, Y6.0, Y7.5, Y10-Y11) and the BRACO-LD cohort.

DNA extraction from buccal swabs was performed using the QIAsymphony SP (QSSP) automated extractor (Qiagen, Hillden, Germany) with QSSP DSP DNA midi kit according to manufacturer's protocol. Extracted DNA was used for DNA methylation profiling and/or telomere length analysis (depending on the quantity of DNA extracted).

Epigenetic Age (PedBE): Genomic DNA was bisulfite converted using the EZ-96 DNA Methylation-Gold Kit (Zymo Research, CA, USA). DNA methylation profiling was performed using the Infinium MethylationEPIC v2.0 BeadChip array following standard protocol (Illumina, CA, USA). DNA methylation .idat files were imported into R[18] for data preprocessing and quality control using the minfi package[19]. Briefly, probes with fewer than three beads for either the methylated or unmethylated channel, or with detection $p \geq 0.01$ were removed. The percent methylation in beta value was normalized using the preprocessNoob method for computing DNA methylation age. 21 Samples with minfi QC score < 9.5 and colMeans(detection $p \geq 0.01$) > 0.06 were excluded. The PedBE was computed using the DNAmAge function from the methylclock R package v0.99.25[20]. The PedBE was available for 3 GUSTO time-points (Y4.5, Y7.5, Y10.5, n = 498) and BRACO-LD (n = 93).

Telomere Length: A validated quantitative polymerase chain reaction (qPCR) protocol was used to measure telomere length (TL)[21]. Commercially available human genomic DNA (G3041) was included as an experimental control on each qPCR plate. TL was quantified using the LightCycler 480 instrument. Forward and reverse primers for telomere detection and human β-globin (reference gene) can be found in Supplementary Table 1 in Lau et al.[21]. The relative TL was calculated as the ratio of telomere (T) to β-globin (S) gene copy numbers (T/S ratio). An average normalizing factor was applied to adjust T/S ratio values across samples. All samples were measured in duplicate, and average values were used for analysis. The coefficient of variation within samples was maintained below 2%, and between-plate variation was 9.14%. Telomere length was available for 4 GUSTO time-points (Y4.5, Y6, Y7.5, Y10.5, n = 858) and BRACO-LD (n = 957, n = 99).

**BrainAge: Model training**

The GPBoost algorithm from the gpboost package v1.5.8[22] was used to train BrainAge models predicting chronological age from neuroimaging features with a random effects term for participant ID. GPBoost combines tree-boosting, Gaussian processes with mixed-effects models, allowing for the implementation of machine learning models in longitudinal data[23].

We employed 5-fold repeated cross-validation with 5 repeats to optimize model hyperparameters within the Train dataset. Partitioning was grouped by participant ID to ensure that all scans from an individual remained within the same internal training or validation set. In each iteration, 20% of the Train participants were held out for validation, and optimal hyperparameters were selected by minimizing Root Mean Square Error (RMSE). Optimization was conducted using the gpboost gpb.grid.search.tune.parameters function with the following hyperparameter tuning grid (360 combinations):

- Learning Rate ∈ {0.01, 0.1, 1}
- Max Tree Depth ∈ {1, 2, 3, 5, 10, 15)
- Min Data in Leaf ∈ {5, 10, 50,100}
- $L_2$ Lambda ∈ {0, 1, 10, 25, 50}

Number of boosting iterations was set at 2000 with an early stopping round of 10.

The final model was then retrained with the selected hyperparameters on the full Train dataset.

**Subgroup Comparisons: Details**

Participants were categorized into subgroups based on a range of outcomes (Comparison 1 & 2) and exposures (Comparison 3 & 4) covering bother the early- and late-childhood phases.

Comparison 1: BRACO-LD vs GUSTO/S-PRESTO

$$BrainAge \sim \beta_0 + \beta_1 BRACO - LD + \beta_2 Age$$

BRACO-LD was coded as 1 for participants in the BRACO-LD cohort and 0 for participants from GUSTO and S-PRESTO. BrainAge estimates were restricted to the early-childhood phase to match the BRACO-LD age range (Y4-8), corresponding to the Y4.5, Y6, and Y7.5 time-points in GUSTO. 1 time-point was randomly selected for participants with longitudinal data. The final sample numbers:

| BRACO-LD = 1 | BRACO-LD = 0 | | | |
|---|---|---|---|---|
| BRACO-LD | GUSTO Y4.5 | GUSTO Y6 | GUSTO Y7.5 | S-PRESTO |
| 214 | 36 | 36 | 53 | 20 |

$\beta_1$ represents the magnitude and direction in which BrainAge estimates in BRACO-LD differs from that of GUSTO and S-PRESTO participants.

Comparison 2: Youth Self Report Total Problems (Sex-stratified)

$$BrainAge \sim \beta_0 + \beta_1 High\_YSR + \beta_2 Age$$

High YSR was coded as 1 for participants who scored > 63 for YSR[24] total problems T score and 0 for participants who scored ≤ 63 for YSR total problems T score. BrainAge estimates at GUSTO Y13 were used. The final sample numbers:

| Females | | Males | |
|---|---|---|---|
| High Stress =1 | High Stress = 0 | High Stress =1 | High Stress = 0 |
| 10 | 27 | 11 | 33 |

$\beta_1$ represents the magnitude and direction in which BrainAge estimates in the High YSR subgroup differs from individuals that don't meet the cut-off.

Comparison 3: Cumulative ELA

$$BrainAge \sim \beta_0 + \beta_1 ELA + \beta_2 Age$$

ELA was scored based on how many ELA components they met criteria for, generating an ordinal cumulative score ranging from 0 (meeting criteria for 0 categories) to 5 (met criteria for 5 components). BrainAge estimates were restricted to the early-childhood phase for the GUSTO cohort. 1 time-point was randomly selected for participants with longitudinal data. The final sample numbers:

| ELA = 0 | ELA = 1 | ELA = 2 | ELA = 3 | ELA = 4 | ELA = 5 |
|---|---|---|---|---|---|
| 15 | 15 | 24 | 27 | 9 | 3 |

$\beta_1$ represents the change in BrainAge estimates for each additional level of ELA.

Comparison 4: Perceived Stress Scale (Sex-stratified)

$$BrainAge \sim \beta_0 + \beta_1 High\_Stress + \beta_2 Age$$

High Stress was coded as 1 for participants who scored > 21 for PSS total score (corresponding to the top 10 percentile of the full PSS cohort) and 0 for participants who scored ≤ 21 for PSS total score. BrainAge estimates at GUSTO Y10.5 were used. The final sample numbers:

| Females | | Males | |
|---|---|---|---|
| High Stress =1 | High Stress = 0 | High Stress =1 | High Stress = 0 |
| 8 | 32 | 7 | 48 |

$\beta_1$ represents the magnitude and direction in which BrainAge estimates in the High Stress subgroup differs from individuals that don't meet the cut-off.

Supplementary Fig S1: Study flowchart and analysis numbers

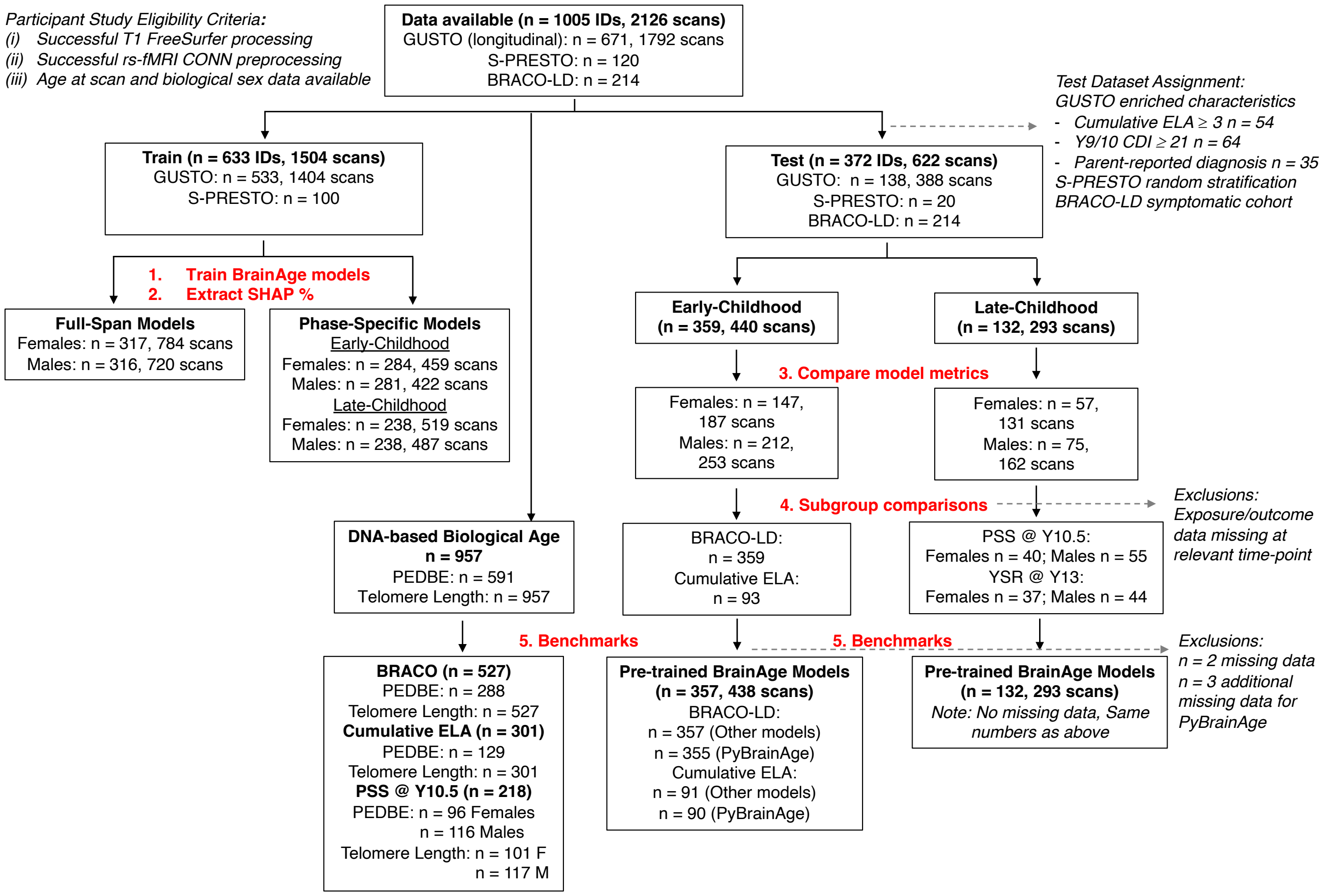


Fig S1: Study data were from three Singaporean pediatric cohorts (age range: 4 to 13 years). Participants were included if their T1 scan and resting state fMRI scan successfully went through pre-processing at at least 1 time-point. Included participants also had data available on age at scan and biological sex. A subset of participants also had DNA-based biological age measures (PedBE epigenetic clock, telomere length) calculated. Participants were then split into a Train and Test dataset. BrainAge models were trained in the Train dataset. BrainAge estimates were then calculated in the Test dataset. The sample size for subgroup comparisons was determined by the participants in the Test dataset who also had data for the instrument used to categorize subgroups.
Note: CDI, Child Depression Inventory 2nd version; ELA, early life adversity; PedBE, Pediatric-Buccal-Epigenetic; PSS, Perceived Stress Scale; YSR, Youth Self Report

Supplementary Fig S2: Sex-stratified regression estimates for BRACO and cumulative early life adversity

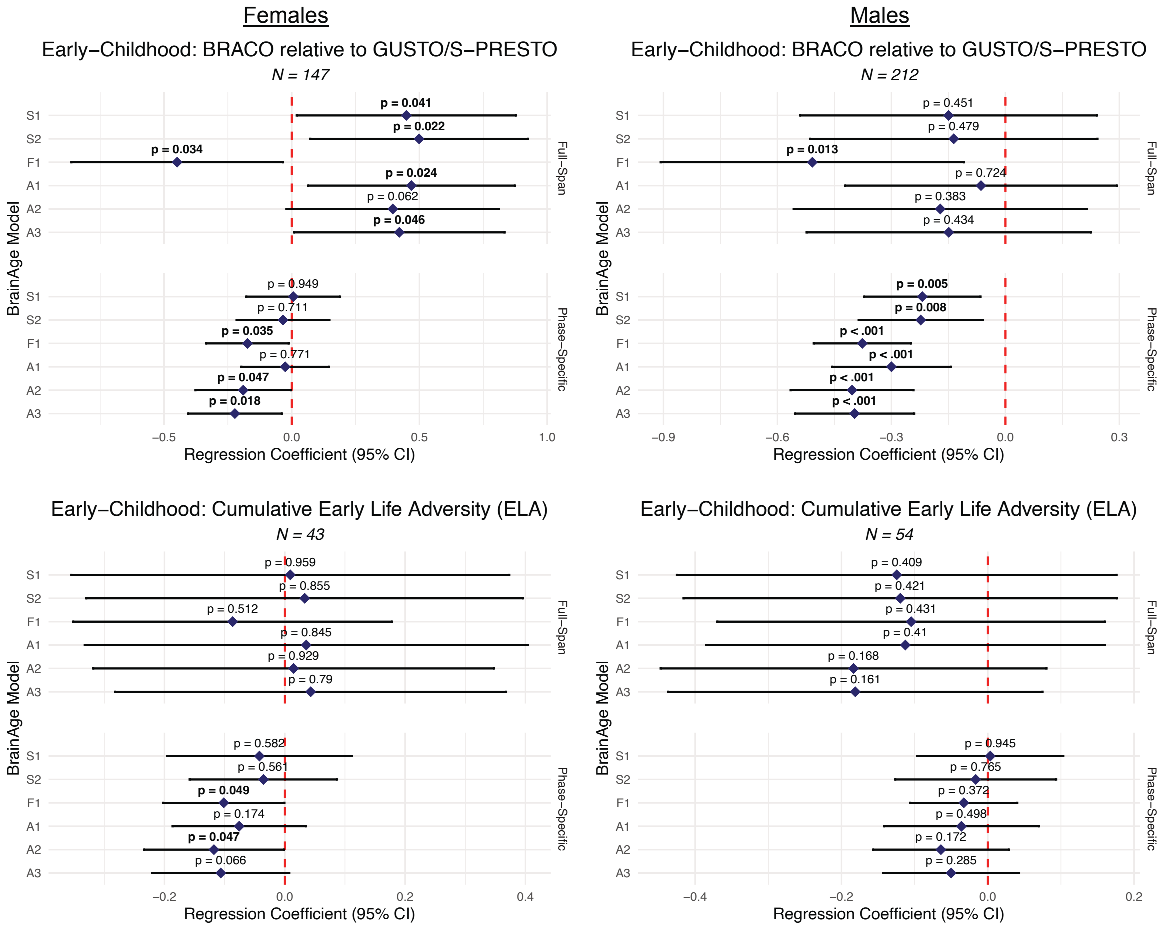


Fig S2: Sex-stratified estimated difference in BrainAge for the following symptomatic subgroups: (i) symptomatic BRACO-LD cohort and (ii) cumulative early life adversity. Significant regression estimates are largely in the same direction in males and females and full (combined males and females) cohort estimates are reported in the main text.

Supplementary Fig S3: Partial Pearson's correlations between BrainAge and biological age measures

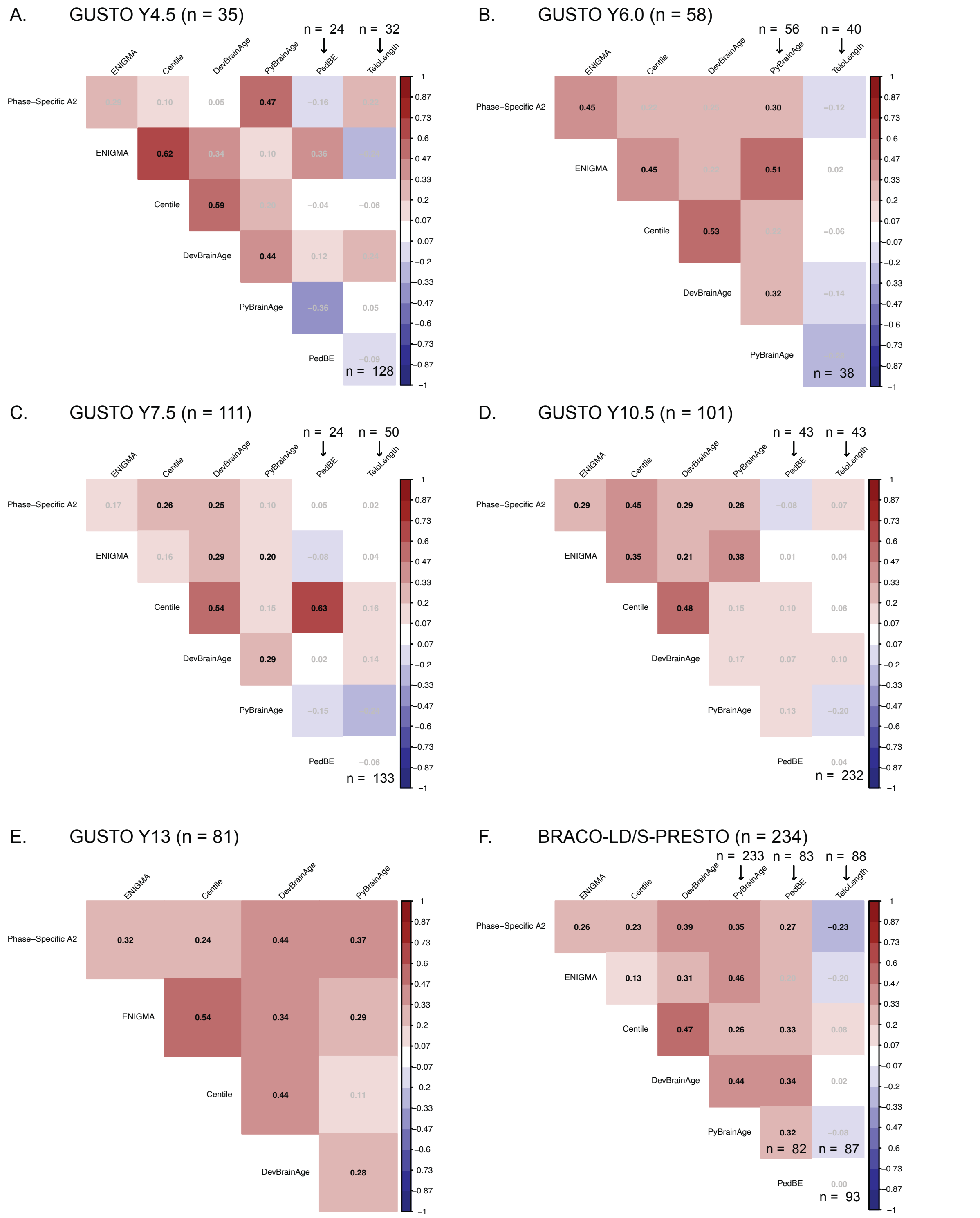


Fig S3: Partial Pearson's correlations between BrainAge and biological age measures controlling for sex. Each correlation matrix consisted of age measures available at that time-point. For example, the PedBE is not available at GUSTO Y6. DNA-based age measures were not available at age 13 years and in the S-PRESTO cohort. Sample size is stated in the plot title unless otherwise stated in the specific cell or column (downward arrow). Correlation values that do not pass significance ($p < 0.05$) are displayed in grey text.

Supplementary Fig S4a-b: Sensitivity Analyses for MRI Motion and Harmonization

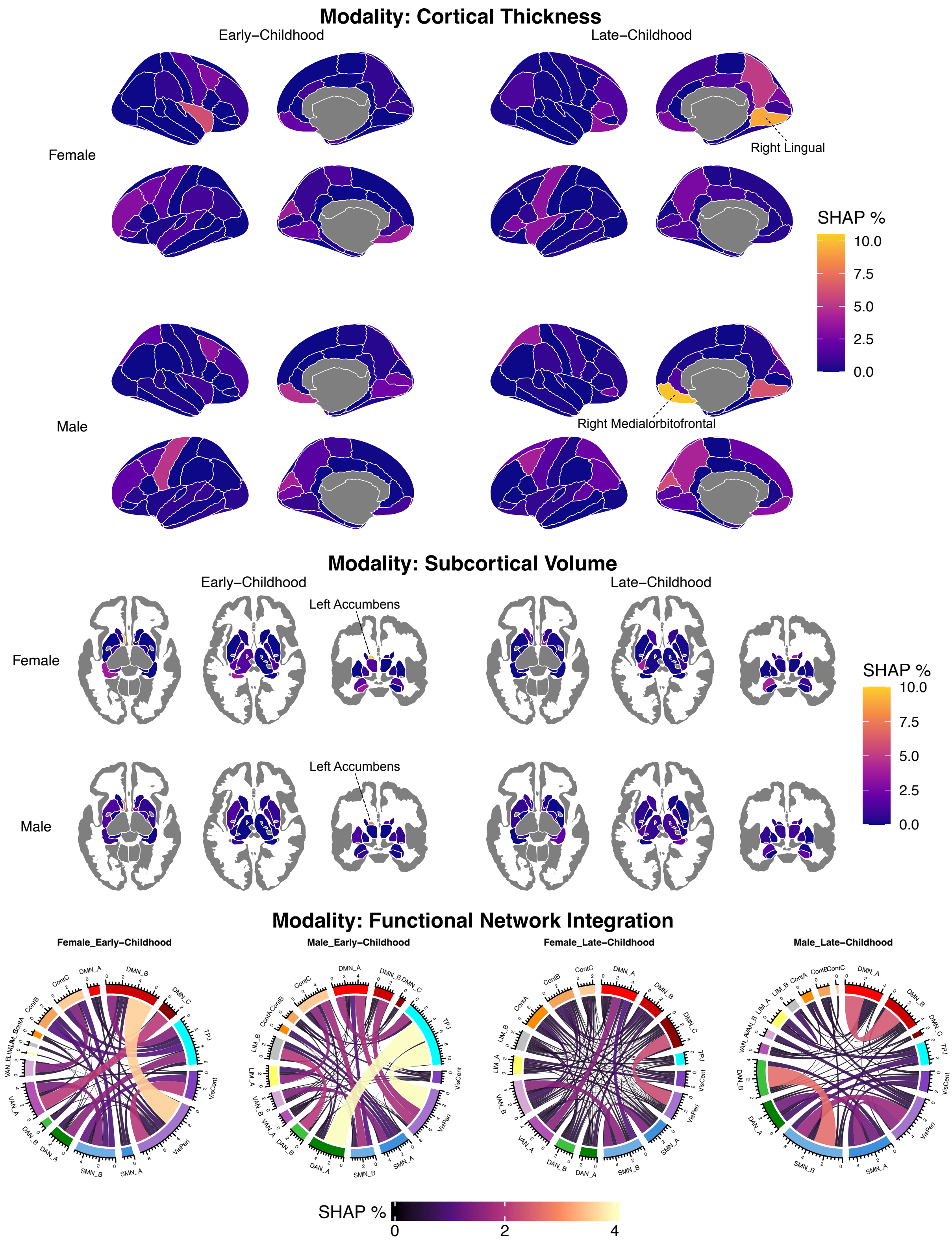


Fig S4a: SHAP values for each neuroimaging feature (split by modality) displayed as a percentage of total SHAP values for the Phase-Specific A2 (M) model that is trained only on scans that passed MRI motion criteria. We observed similar SHAP patterns to the main analysis, with different patterns for the Early-Childhood and Late-Childhood models.

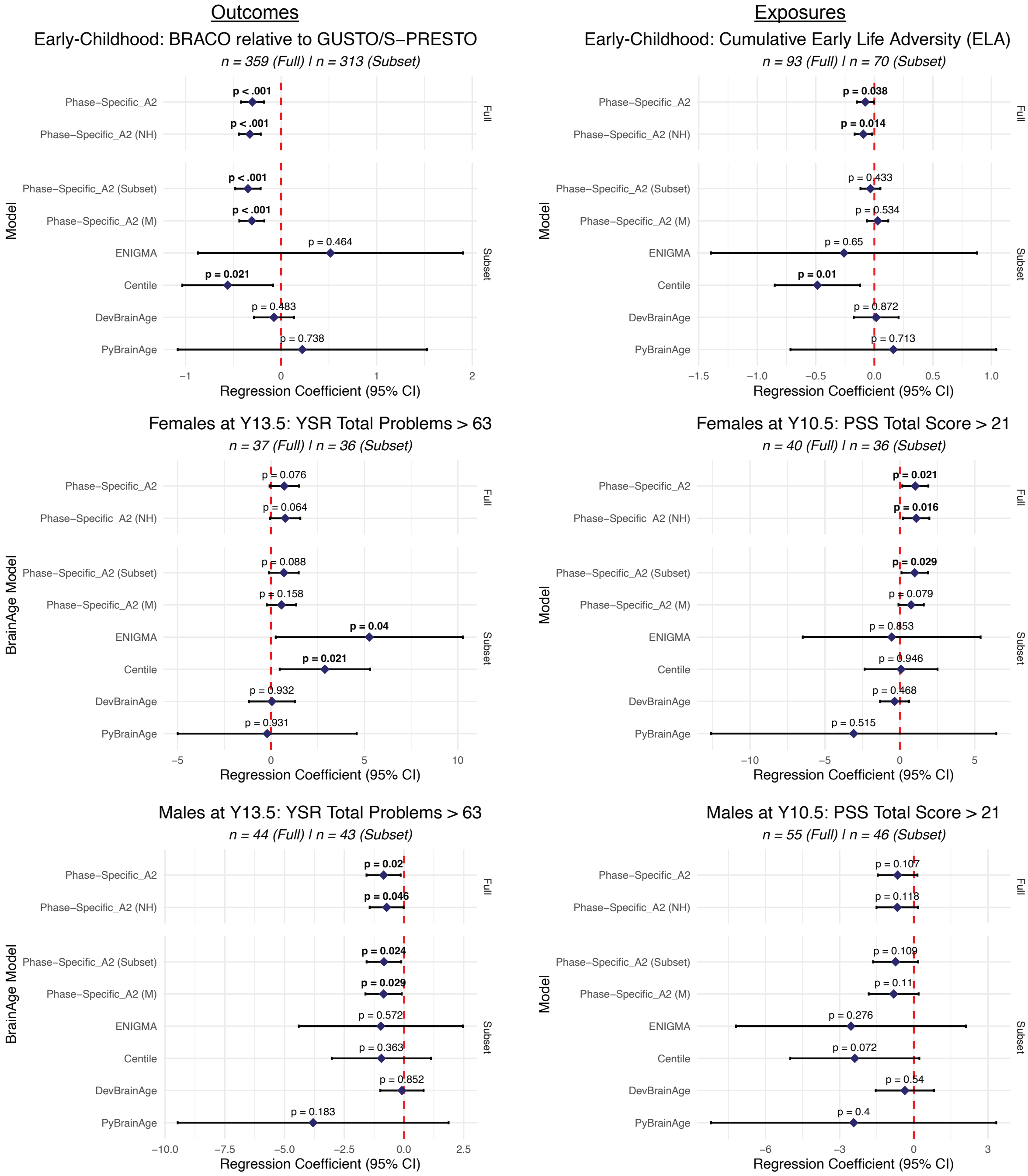


Fig S4b: Full data consists of the main Phase-Specific A2 model and the model applied to non-harmonized testing data (Phase-Specific_A2 (NH)). Subset data consist of scans that passed the motion criteria. Phase-Specific A2 (Subset) is the main model applied to the subset of scans that passed motion criteria, Phase-Specific A2 (M) is the model retrained on the subset. Regression coefficient estimates are similar among all 4 versions, although the p-value is affected for cumulative ELA especially, possibly due to lower statistical power due to smaller sample size. Results suggest our findings are robust to harmonization and motion.

Supplementary Tables S1a-b: Summary of demographic and MRI QC measures by cohort and time-points

Table S1a: Summary of demographics by cohort

| Characteristic | Overall | GUSTO | S-PRESTO | BRACO-LD | p |
|---|---|---|---|---|---|
| n | 1005 | 671 | 120 | 214 | |
| Sex = Male (%) | 538 (53.5) | 339 (50.5) | 61 (50.8) | 138 ( 64.5) | 0.001 |
| Ethnicity (%) | | | | | <0.001 |
| Chinese | 589 (58.6) | 365 (54.4) | 95 (79.2) | 129 ( 60.3) | |
| Indian | 133 (13.2) | 114 (17.0) | 7 ( 5.8) | 12 ( 5.6) | |
| Malay | 261 (26.0) | 191 (28.5) | 16 (13.3) | 54 ( 25.2) | |
| Other | 22 ( 2.2) | 1 ( 0.1) | 2 ( 1.7) | 19 ( 8.9) | |
| Dataset = Test (%) | 372 (37.0) | 138 (20.6) | 20 (16.7) | 214 (100.0) | <0.001 |

Table S1b: Summary of Age and MRI QC measures by cohort and time-point

| Timepoint | Age in years (Range) | Scans (n) | Mean Euler (SD) | Mean fMRI Motion (SD) |
|---|---|---|---|---|
| GUSTO | | | | |
| Y4.5 | 4.58 (4.44-4.94) | 204 | -119.373 (104.016) | 0.149 (0.087) |
| Y6.0 | 6.04 (5.82-6.6) | 289 | -100.291 (81.701) | 0.142 (0.08) |
| Y7.5 | 7.46 (7.25-8.03) | 494 | -68.599 (43.646) | 0.14 (0.087) |
| Y10.5 | 10.74 (10.38-11.35) | 466 | -49.545 (53.308) | 0.165 (0.085) |
| Y13 | 12.99 (12.62-13.7) | 339 | -42.336 (26.932) | 0.133 (0.065) |
| S-PRESTO | | | | |
| Y4-6 | 5.27 (4.57-7.31) | 120 | -70.6 (54.738) | 0.199 (0.049) |
| BRACO-LD | | | | |
| Y4-8 | 5.84 (4-7.99) | 214 | -74.757 (66.922) | 0.188 (0.056) |

Supplementary Table S2: Age and MRI QC measures stratified by Train/Test and time-point

<u>Table S2: Summary of Age and MRI QC measures by Train/Test dataset and time-point</u>

| | | | Train Dataset | | Test Dataset | | Comparison[a] |
|---|---|---|---|---|---|---|---|
| Variable | Cohort | Timepoint | n | Mean (SD) | n | Mean (SD) | p |
| Age in years | GUSTO | Y4.5 | 168 | 4.59 (0.09) | 36 | 4.57 (0.07) | 0.578 |
| | | Y6.0 | 230 | 6.05 (0.12) | 59 | 6.02 (0.13) | 0.094 |
| | | Y7.5 | 383 | 7.47 (0.15) | 111 | 7.44 (0.14) | 0.042 |
| | | Y10.5 | 365 | 10.74 (0.19) | 101 | 10.73 (0.20) | 0.443 |
| | | Y13 | 258 | 12.98 (0.18) | 81 | 13.02 (0.21) | 0.176 |
| | S-PRESTO | Y4-6 | 100 | 5.24 (0.49) | 20 | 5.38 (0.67) | 0.526 |
| | BRACO-LD | Y4-8 | 0 | NaN (NA) | 214 | 5.84 (0.97) | |
| Mean Euler | GUSTO | Y4.5 | 168 | -119.37 (100.35) | 36 | -119.39 (121.24) | 0.351 |
| | | Y6.0 | 230 | -98.44 (74.14) | 59 | -107.49 (106.66) | 0.405 |
| | | Y7.5 | 383 | -68.33 (44.25) | 111 | -69.53 (41.67) | 0.889 |
| | | Y10.5 | 365 | -49.66 (55.34) | 101 | -49.13 (45.47) | 0.754 |
| | | Y13 | 258 | -40.78 (19.82) | 81 | -47.31 (42.06) | 0.893 |
| | S-PRESTO | Y4-6 | 100 | -73.66 (58.87) | 20 | -55.30 (20.26) | 0.360 |
| | BRACO-LD | Y4-8 | 0 | NaN (NA) | 214 | -74.76 (66.92) | |
| Mean fMRI Motion | GUSTO | Y4.5 | 168 | 0.15 (0.09) | 36 | 0.16 (0.08) | 0.350 |
| | | Y6.0 | 230 | 0.14 (0.08) | 59 | 0.14 (0.08) | 0.981 |
| | | Y7.5 | 383 | 0.14 (0.09) | 111 | 0.14 (0.09) | 0.837 |
| | | Y10.5 | 365 | 0.16 (0.08) | 101 | 0.17 (0.09) | 0.470 |
| | | Y13 | 258 | 0.14 (0.07) | 81 | 0.13 (0.06) | 0.379 |
| | S-PRESTO | Y4-6 | 100 | 0.20 (0.05) | 20 | 0.20 (0.06) | 0.852 |
| | BRACO-LD | Y4-8 | 0 | NaN (NA) | 214 | 0.19 (0.06) | |

[a]Wilcox Test performed for non-parametric group comparisons between Train and Test Datasets

Supplementary Tables S3a-f: Regression estimates across candidate BrainAge models

Table S3a: Summary of Regression Results for BRACO

| Type | Feature | β estimate [95% CI] | Std Error | t statistic | p | sig |
|---|---|---|---|---|---|---|
| Full-Span | S1 | 0.130 [-0.16, 0.42] | 0.146 | 0.894 | 0.372 | N.S |
| | S2 | 0.156 [-0.12, 0.44] | 0.143 | 1.095 | 0.274 | N.S |
| | F1 | -0.471 [-0.76, -0.19] | 0.145 | -3.257 | 0.001 | ** |
| | A1 | 0.179 [-0.09, 0.44] | 0.135 | 1.322 | 0.187 | N.S |
| | A2 | 0.069 [-0.21, 0.35] | 0.143 | 0.486 | 0.628 | N.S |
| | A3 | 0.093 [-0.18, 0.37] | 0.140 | 0.670 | 0.504 | N.S |
| Phase-Specific | S1 | -0.115 [-0.23, 0.00] | 0.059 | -1.942 | 0.053 | . |
| | S2 | -0.137 [-0.26, -0.02] | 0.061 | -2.242 | 0.026 | * |
| | F1 | -0.279 [-0.38, -0.18] | 0.051 | -5.493 | $7.5 \times 10^{-08}$ | *** |
| | A1 | -0.180 [-0.29, -0.07] | 0.058 | -3.085 | 0.002 | ** |
| | A2 | -0.300 [-0.42, -0.18] | 0.061 | -4.886 | $1.6 \times 10^{-06}$ | *** |
| | A3 | -0.308 [-0.43, -0.19] | 0.060 | -5.136 | $4.6 \times 10^{-07}$ | *** |

Table S3b: Summary of Regression Results for High YSR Total (Females)

| Type | Feature | β estimate [95% CI] | Std Error | t statistic | p | sig |
|---|---|---|---|---|---|---|
| Full-Span | S1 | 0.403 [-0.55, 1.35] | 0.468 | 0.860 | 0.396 | N.S. |
| | S2 | 0.667 [-0.21, 1.55] | 0.434 | 1.538 | 0.133 | N.S. |
| | F1 | -0.072 [-1.27, 1.13] | 0.590 | -0.123 | 0.903 | N.S. |
| | A1 | 0.698 [-0.15, 1.55] | 0.418 | 1.669 | 0.104 | N.S. |
| | A2 | 0.561 [-0.23, 1.35] | 0.390 | 1.439 | 0.159 | N.S. |
| | A3 | 0.539 [-0.26, 1.34] | 0.392 | 1.377 | 0.177 | N.S. |
| Phase-Specific | S1 | 0.351 [-0.55, 1.25] | 0.441 | 0.796 | 0.431 | N.S. |
| | S2 | 0.532 [-0.22, 1.29] | 0.372 | 1.430 | 0.162 | N.S. |
| | F1 | -0.032 [-0.91, 0.85] | 0.433 | -0.074 | 0.942 | N.S. |
| | A1 | 0.490 [-0.25, 1.23] | 0.366 | 1.341 | 0.189 | N.S. |
| | A2 | 0.707 [-0.08, 1.49] | 0.386 | 1.833 | 0.076 | . |
| | A3 | 0.488 [-0.18, 1.15] | 0.328 | 1.488 | 0.146 | N.S. |

Table S3c: Summary of Regression Results for High YSR Total (Males)

| Type | Feature | β estimate [95% CI] | Std Error | t statistic | p | sig |
|---|---|---|---|---|---|---|
| Full-Span | S1 | -0.610 [-1.57, 0.35] | 0.474 | -1.286 | 0.206 | N.S. |
| | S2 | -0.718 [-1.65, 0.22] | 0.463 | -1.552 | 0.128 | N.S. |
| | F1 | 0.297 [-0.54, 1.14] | 0.417 | 0.713 | 0.480 | N.S. |
| | A1 | -0.681 [-1.60, 0.24] | 0.456 | -1.493 | 0.143 | N.S. |
| | A2 | -0.747 [-1.68, 0.18] | 0.460 | -1.624 | 0.112 | N.S. |
| | A3 | -0.763 [-1.68, 0.16] | 0.456 | -1.674 | 0.102 | N.S. |
| Phase-Specific | S1 | -0.386 [-1.11, 0.34] | 0.358 | -1.079 | 0.287 | N.S. |
| | S2 | -0.638 [-1.39, 0.11] | 0.372 | -1.718 | 0.093 | . |
| | F1 | -0.099 [-0.61, 0.41] | 0.251 | -0.393 | 0.697 | N.S. |
| | A1 | -0.433 [-1.20, 0.34] | 0.380 | -1.138 | 0.262 | N.S. |
| | A2 | -0.853 [-1.57, -0.14] | 0.354 | -2.412 | 0.020 | * |
| | A3 | -0.821 [-1.52, -0.12] | 0.348 | -2.356 | 0.023 | * |

Table S3d: Summary of Regression Results for Cumulative ELA

| Type | Feature | β estimate [95% CI] | Std Error | t statistic | p | sig |
|---|---|---|---|---|---|---|
| Full-Span | S1 | -0.087 [-0.31, 0.14] | 0.112 | -0.775 | 0.440 | N.S. |
| | S2 | -0.089 [-0.31, 0.13] | 0.112 | -0.791 | 0.431 | N.S. |
| | F1 | -0.052 [-0.24, 0.13] | 0.093 | -0.564 | 0.574 | N.S. |
| | A1 | -0.079 [-0.29, 0.14] | 0.108 | -0.729 | 0.468 | N.S. |
| | A2 | -0.116 [-0.32, 0.09] | 0.101 | -1.145 | 0.255 | N.S. |
| | A3 | -0.098 [-0.29, 0.10] | 0.099 | -0.999 | 0.321 | N.S. |
| Phase-Specific | S1 | -0.029 [-0.11, 0.06] | 0.043 | -0.674 | 0.502 | N.S. |
| | S2 | -0.025 [-0.10, 0.05] | 0.039 | -0.641 | 0.523 | N.S. |
| | F1 | -0.050 [-0.11, 0.01] | 0.031 | -1.621 | 0.109 | N.S. |
| | A1 | -0.063 [-0.14, 0.01] | 0.038 | -1.656 | 0.101 | N.S. |
| | A2 | -0.077 [-0.15, -0.00] | 0.037 | -2.102 | 0.038 | * |
| | A3 | -0.067 [-0.14, 0.01] | 0.036 | -1.840 | 0.069 | . |

Table S3e: Summary of Regression Results for High PSS (Females)

| Type | Feature | β estimate [95% CI] | Std Error | t statistic | p | sig |
|---|---|---|---|---|---|---|
| Full-Span | S1 | 0.488 [-0.92, 1.90] | 0.695 | 0.702 | 0.487 | N.S. |
| | S2 | 1.033 [-0.20, 2.27] | 0.610 | 1.694 | 0.099 | . |
| | F1 | 0.449 [-0.42, 1.32] | 0.429 | 1.046 | 0.302 | N.S. |
| | A1 | 0.938 [-0.37, 2.25] | 0.646 | 1.451 | 0.155 | N.S. |
| | A2 | 0.814 [-0.46, 2.09] | 0.631 | 1.291 | 0.205 | N.S. |
| | A3 | 0.737 [-0.51, 1.99] | 0.617 | 1.194 | 0.240 | N.S. |
| Phase-Specific | S1 | 0.322 [-0.70, 1.34] | 0.502 | 0.642 | 0.525 | N.S. |
| | S2 | 0.547 [-0.34, 1.43] | 0.436 | 1.252 | 0.218 | N.S. |
| | F1 | 0.237 [-0.38, 0.86] | 0.307 | 0.774 | 0.444 | N.S. |
| | A1 | 0.549 [-0.31, 1.40] | 0.422 | 1.302 | 0.201 | N.S. |
| | A2 | 1.033 [0.16, 1.90] | 0.429 | 2.408 | 0.021 | * |
| | A3 | 0.665 [-0.22, 1.55] | 0.438 | 1.518 | 0.137 | N.S. |

Table S3f: Summary of Regression Results for High PSS (Males)

| Type | Feature | β estimate [95% CI] | Std Error | t statistic | p | sig |
|---|---|---|---|---|---|---|
| Full-Span | S1 | -0.910 [-2.11, 0.29] | 0.599 | -1.518 | 0.135 | N.S. |
| | S2 | -0.837 [-2.03, 0.36] | 0.595 | -1.408 | 0.165 | N.S. |
| | F1 | 0.077 [-0.95, 1.10] | 0.511 | 0.151 | 0.881 | N.S. |
| | A1 | -0.805 [-2.02, 0.41] | 0.608 | -1.325 | 0.191 | N.S. |
| | A2 | -1.125 [-2.34, 0.09] | 0.604 | -1.864 | 0.068 | . |
| | A3 | -1.026 [-2.26, 0.21] | 0.615 | -1.668 | 0.101 | N.S. |
| Phase-Specific | S1 | -0.836 [-1.65, -0.02] | 0.405 | -2.065 | 0.044 | * |
| | S2 | -0.797 [-1.59, -0.00] | 0.395 | -2.018 | 0.049 | * |
| | F1 | 0.107 [-0.55, 0.77] | 0.329 | 0.325 | 0.746 | N.S. |
| | A1 | -0.930 [-1.63, -0.23] | 0.348 | -2.676 | 0.010 | ** |
| | A2 | -0.655 [-1.46, 0.15] | 0.400 | -1.639 | 0.107 | N.S. |
| | A3 | -0.676 [-1.47, 0.12] | 0.396 | -1.710 | 0.093 | . |

Supplementary Table S4: Top contributing regions based on SHAP values for selected model Phase-Specific A2

| | Female: Early-childhood | | Male: Early-childhood | | Female: Late-childhood | | Male: Late-childhood | |
|---|---|---|---|---|---|---|---|---|
| | Region | SHAP (%) | Region | SHAP (%) | Region | SHAP (%) | Region | SHAP (%) |
| 1 | left_accumbens_area | 9.11 | left_accumbens_area | 6.59 | rh_lingual_thickness | 7.6 | rh_medialorbitofrontal_thickness | 8.9 |
| 2 | rh_frontalpole_thickness | 4.09 | lh_cuneus_thickness | 4.73 | rh_precuneus_thickness | 4.37 | lh_cuneus_thickness | 5.25 |
| 3 | rh_insula_thickness | 3.85 | lh_precentral_thickness | 4.51 | left_amygdala | 4.1 | rh_lingual_thickness | 4.42 |
| 4 | Int.VisPerixDMN_B | 3.36 | right_accumbens_area | 4.23 | lh_insula_thickness | 3.18 | lh_precuneus_thickness | 4.4 |
| 5 | lh_cuneus_thickness | 2.97 | rh_medialorbitofrontal_thickness | 3.82 | right_accumbens_area | 3.01 | rh_cuneus_thickness | 4.06 |
| 6 | rh_medialorbitofrontal_thickness | 2.8 | Int.VisPerixTPJ | 3.69 | rh_medialorbitofrontal_thickness | 2.41 | rh_superiorparietal_thickness | 3.89 |
| 7 | lh_lingual_thickness | 2.51 | Int.DAN_AxTPJ | 3.67 | rh_parstriangularis_thickness | 2.24 | lh_superiorfrontal_thickness | 3.14 |
| 8 | Int.VAN_AxDMN_C | 2.45 | rh_pericalcarine_thickness | 3.53 | lh_precentral_thickness | 2.15 | right_accumbens_area | 2.94 |
| 9 | modularity | 2.35 | rh_frontalpole_thickness | 3.37 | rh_cuneus_thickness | 2.12 | lh_medialorbitofrontal_thickness | 2.77 |
| 10 | lh_medialorbitofrontal_thickness | 2.12 | rh_caudalmiddlefrontal_thickness | 3.05 | rh_lateralorbitofrontal_thickness | 1.77 | lh_pericalcarine_thickness | 2.71 |
| 11 | Int.VisCentxVisPeri | 2.07 | rh_superiorparietal_thickness | 2.78 | lh_precuneus_thickness | 1.76 | lh_insula_thickness | 2.33 |
| 12 | Int.VAN_BxContC | 2.05 | lh_rostralmiddlefrontal_thickness | 2.14 | lh_parstriangularis_thickness | 1.65 | left_amygdala | 2.09 |
| 13 | lh_caudalmiddlefrontal_thickness | 2.05 | rh_rostralmiddlefrontal_thickness | 2.01 | rh_isthmuscingulate_thickness | 1.48 | lh_lingual_thickness | 2.03 |
| 14 | lh_rostralmiddlefrontal_thickness | 1.96 | lh_paracentral_thickness | 1.84 | lh_lingual_thickness | 1.39 | Int.SMN_BxDAN_B | 1.91 |
| 15 | lh_rostralanteriorcingulate_thickness | 1.95 | Int.SMN_BxContC | 1.71 | Int.VisPerixDMN_C | 1.35 | lh_caudalmiddlefrontal_thickness | 1.84 |
| 16 | rh_caudalmiddlefrontal_thickness | 1.88 | lh_pericalcarine_thickness | 1.66 | lh_bankssts_thickness | 1.27 | rh_precentral_thickness | 1.69 |
| 17 | Int.DAN_AxTPJ | 1.81 | rh_insula_thickness | 1.37 | rh_posteriorcingulate_thickness | 1.24 | Int.SMN_AxSMN_B | 1.59 |
| 18 | lh_precentral_thickness | 1.81 | lh_frontalpole_thickness | 1.23 | rh_superiorfrontal_thickness | 1.24 | right_thalamus_proper | 1.59 |
| 19 | rh_entorhinal_thickness | 1.47 | Int.ContBxDMN_C | 1.2 | right_caudate | 1.13 | lh_precentral_thickness | 1.55 |
| 20 | rh_rostralmiddlefrontal_thickness | 1.43 | rh_superiortemporal_thickness | 1.14 | Int.SMN_BxVAN_B | 1.11 | Int.VisCentxSMN_B | 1.53 |
| 21 | left_hippocampus | 1.41 | Int.VisPerixVAN_B | 1.14 | lh_isthmuscingulate_thickness | 1.04 | Int.VisPerixSMN_A | 1.33 |
| 22 | Int.VisPerixVAN_B | 1.32 | Int.VisPerixDMN_B | 1.07 | rh_inferiorparietal_thickness | 1.02 | lh_bankssts_thickness | 1.33 |
| 23 | left_amygdala | 1.25 | Int.ContCxTPJ | 0.97 | lh_posteriorcingulate_thickness | 0.91 | right_hippocampus | 1.28 |
| 24 | rh_lingual_thickness | 1.23 | lh_medialorbitofrontal_thickness | 0.9 | Int.SMN_BxDMN_A | 0.9 | Int.VisPerixSMN_B | 1.17 |
| 25 | Int.ContBxTPJ | 1.23 | lh_superiortemporal_thickness | 0.89 | lh_pericalcarine_thickness | 0.9 | lh_inferiorparietal_thickness | 1.12 |
| 26 | left_thalamus_proper | 1.21 | right_pallidum | 0.85 | Int.SMN_AxSMN_B | 0.88 | lh_parsorbitalis_thickness | 1.04 |
| 27 | Int.SMN_BxContC | 1.21 | left_putamen | 0.8 | Int.LIM_AxDMN_A | 0.86 | rh_parsorbitalis_thickness | 1.01 |
| 28 | lh_pericalcarine_thickness | 1.14 | rh_bankssts_thickness | 0.79 | rh_rostralmiddlefrontal_thickness | 0.82 | lh_superiorparietal_thickness | 0.95 |
| 29 | Int.SMN_BxContB | 1.1 | Int.VisPerixLIM_B | 0.76 | Int.VAN_BxContA | 0.72 | right_amygdala | 0.89 |
| 30 | lh_parahippocampal_thickness | 1.05 | rh_entorhinal_thickness | 0.75 | left_caudate | 0.72 | left_caudate | 0.86 |
| 31 | lh_parsorbitalis_thickness | 1.04 | Int.SMN_AxVAN_B | 0.74 | Int.DAN_BxLIM_B | 0.71 | rh_parsopercularis_thickness | 0.8 |
| 32 | lh_entorhinal_thickness | 1.03 | Int.SMN_AxContA | 0.7 | Int.ContAxContB | 0.7 | Int.LIM_BxLIM_A | 0.78 |
| 33 | lh_precuneus_thickness | 0.96 | Int.VAN_AxLIM_A | 0.68 | Int.ContBxDMN_B | 0.7 | rh_rostralanteriorcingulate_thickness | 0.78 |
| 34 | Int.VisPerixLIM_B | 0.93 | Int.VisCentxContA | 0.68 | right_amygdala | 0.67 | right_pallidum | 0.75 |
| 35 | lh_supramarginal_thickness | 0.92 | Int.DAN_BxContC | 0.67 | lh_medialorbitofrontal_thickness | 0.67 | right_caudate | 0.75 |
| 36 | Int.DAN_AxLIM_B | 0.92 | lh_insula_thickness | 0.67 | Int.SMN_AxDMN_A | 0.66 | Int.VAN_AxDMN_A | 0.74 |
| 37 | Int.LIM_AxTPJ | 0.88 | lh_bankssts_thickness | 0.65 | Int.SMN_AxLIM_A | 0.64 | lh_lateralorbitofrontal_thickness | 0.7 |
| 38 | rh_precentral_thickness | 0.84 | Int.SMN_BxContB | 0.64 | Int.SMN_AxContA | 0.62 | lh_entorhinal_thickness | 0.67 |
| 39 | Int.DMN_BxTPJ | 0.77 | Int.DAN_AxContC | 0.64 | Int.SMN_BxContB | 0.6 | rh_transversetemporal_thickness | 0.66 |
| 40 | left_pallidum | 0.74 | rh_inferiorparietal_thickness | 0.63 | lh_superiorfrontal_thickness | 0.6 | rh_pericalcarine_thickness | 0.58 |
| 41 | right_accumbens_area | 0.72 | Int.SMN_BxLIM_A | 0.63 | Int.LIM_BxDMN_A | 0.59 | Int.DMN_AxDMN_B | 0.56 |
| 42 | rh_parsorbitalis_thickness | 0.67 | Int.DMN_AxTPJ | 0.62 | Int.VAN_AxDMN_A | 0.58 | rh_lateraloccipital_thickness | 0.52 |
| 43 | Int.DMN_AxDMN_C | 0.66 | rh_temporalpole_thickness | 0.62 | Int.VAN_BxContB | 0.55 | Int.DAN_BxContB | 0.47 |
| 44 | Int.VAN_AxTPJ | 0.65 | rh_lingual_thickness | 0.6 | Int.VAN_BxDMN_B | 0.54 | rh_rostralmiddlefrontal_thickness | 0.46 |
| 45 | right_amygdala | 0.62 | left_pallidum | 0.58 | lh_superiortemporal_thickness | 0.52 | Int.DAN_BxTPJ | 0.46 |
| 46 | lh_parstriangularis_thickness | 0.6 | Int.VAN_AxDMN_A | 0.53 | Int.LIM_AxDMN_B | 0.52 | | |
| 47 | lh_frontalpole_thickness | 0.59 | right_caudate | 0.52 | Int.VisCentxDMN_C | 0.51 | | |
| 48 | Int.SMN_AxDMN_B | 0.57 | lh_posteriorcingulate_thickness | 0.51 | Int.VisCentxContA | 0.5 | | |
| 49 | right_pallidum | 0.55 | left_hippocampus | 0.51 | Int.SMN_BxVAN_A | 0.5 | | |
| 50 | Int.VAN_AxVAN_B | 0.55 | lh_precuneus_thickness | 0.49 | lh_parsopercularis_thickness | 0.5 | | |
| 51 | Int.ContAxContC | 0.54 | Int.SMN_BxVAN_B | 0.46 | lh_caudalanteriorcingulate_thickness | 0.49 | | |
| 52 | Int.VisCentxSMN_B | 0.51 | Int.VisPerixDAN_A | 0.46 | rh_parsorbitalis_thickness | 0.48 | | |
| 53 | Int.DMN_AxTPJ | 0.49 | | | Int.VAN_AxContC | 0.46 | | |
| 54 | Int.SMN_AxTPJ | 0.48 | | | lh_entorhinal_thickness | 0.46 | | |
| 55 | lh_superiortemporal_thickness | 0.47 | | | | | | |

Note: Regions with SHAP % values greater than average (>0.456) shown

Supplementary Table S5: Details of 4 pre-trained BrainAge models from the literature

| Model | Sample | Age Range | Features | Sex-specific? |
|---|---|---|---|---|
| ENIGMA BrainAge | 19 cohorts<br>Males<br>**Train**: 952 HC<br>**Test**: 927 HC, 986 MDD<br>Females<br>**Train**: 1236 HC controls<br>**Test**: 927 HC, 986 MDD | 18 to 75 years | FreeSurfer Extracted Features from the Desikan-Killiany cortical atlas and aseg subcortical atlas | Yes |
| Centile BrainAge | **Train**: 35683 HCs<br>**Test**: 2101 HCs | 2 Age bins:<br>5 to 40 years<br>40 to 90 years | | Yes |
| DevBrainAge | 6 cohorts<br>**Train**: 1299 HC<br>**Test**: 322 HC, 150 risk-enriched | 9 to 19 years | | No |
| PyBrainAge | 76 sites<br>**Train:** 29175 HCs<br>**Test**: 29174 | 2 to 100 years | FreeSurfer Extracted Features from the Destrieux cortical atlas and aseg subcortical atlas | No |

Supplementary Table S6: Model performance metrics across selected A2 BrainAge model with pre-trained BrainAge models (n = 731)

| Metric | Sex | BrainAge Estimates | | | | | |
|---|---|---|---|---|---|---|---|
| | | Phase-Specific_A2 | Phase-Specific_A2 (NH) | ENIGMA | Centile | DevBrainAge | PyBrainAge |
| Early-Childhood | | | | | | | |
| Median PAD (IQR) | Female | -0.04 (1.27) | 0.01 (1.32) | 12.25 (7.55) | 3.06 (2.57) | 4.22 (1.66) | 5.75 (5.27) |
| MAE (IQR) | Female | 0.64 (0.72) | 0.66 (0.64) | 12.25 (7.48) | 3.09 (2.5) | 4.22 (1.66) | 5.75 (5.27) |
| Age r [95% CI] | Female | 0.59 [0.49, 0.68] | 0.6 [0.5, 0.69] | -0.04 [-0.19, 0.1] | 0.3 [0.17, 0.43] | 0.38 [0.26, 0.5] | 0.26 [0.12, 0.39] |
| Median PAD (IQR) | Male | -0.1 (1.31) | -0.12 (1.23) | 15.78 (7.02) | 2.64 (2.67) | 4.32 (1.74) | 5.75 (4.59) |
| MAE (IQR) | Male | 0.66 (0.72) | 0.64 (0.71) | 15.78 (7.02) | 2.68 (2.51) | 4.32 (1.74) | 5.75 (4.59) |
| Age r [95% CI] | Male | 0.48 [0.38, 0.57] | 0.52 [0.42, 0.61] | 0.06 [-0.06, 0.18] | 0.4 [0.3, 0.5] | 0.28 [0.16, 0.39] | 0.19 [0.07, 0.31] |
| Late-Childhood | | | | | | | |
| Median PAD (IQR) | Female | 0.1 (1.97) | -0.21 (1.74) | 11.87 (8.03) | 3.32 (2.92) | 2.57 (1.94) | 10.15 (7.53) |
| MAE (IQR) | Female | 0.96 (1.05) | 0.92 (1.14) | 11.87 (7.95) | 3.32 (2.87) | 2.57 (1.88) | 10.15 (7.53) |
| Age r [95% CI] | Female | 0.79 [0.72, 0.85] | 0.79 [0.72, 0.85] | 0.4 [0.24, 0.53] | 0.71 [0.61, 0.78] | 0.75 [0.67, 0.82] | 0.51 [0.37, 0.63] |
| Median PAD (IQR) | Male | 0.09 (1.95) | 0.06 (1.99) | 13.08 (6.92) | 2.28 (2.95) | 2.4 (2.01) | 7.2 (7.94) |
| MAE (IQR) | Male | 0.99 (1.04) | 0.99 (1.03) | 13.08 (6.92) | 2.39 (2.57) | 2.4 (1.99) | 7.2 (7.94) |
| Age r [95% CI] | Male | 0.74 [0.66, 0.8] | 0.77 [0.69, 0.82] | 0.34 [0.2, 0.47] | 0.58 [0.47, 0.68] | 0.74 [0.67, 0.81] | 0.55 [0.43, 0.65] |

Note: Phase-Specific_A2 (NH), Metrics for Phase-Specific A2 model applied to non-harmonized Test dataset

Supplementary Tables S7a-f: Regression estimates for selected A2 BrainAge model and published pre-trained BrainAge and biological age measures

Table S7a: Summary of Regression Results for BRACO

| Model | β estimate [95% CI] | Std Error | t statistic | p | sig |
|---|---|---|---|---|---|
| Phase-Specific_A2 | -0.300 [-0.42, -0.18] | 0.062 | -4.842 | $1.9 \times 10^{-06}$ | *** |
| ENIGMA | 0.284 [-1.01, 1.58] | 0.657 | 0.433 | 0.666 | N.S. |
| Centile | -0.709 [-1.17, -0.25] | 0.233 | -3.039 | 0.003 | ** |
| DevBrainAge | -0.187 [-0.39, 0.01] | 0.103 | -1.822 | 0.069 | . |
| PyBrainAge | 0.357 [-0.80, 1.52] | 0.589 | 0.606 | 0.545 | N.S. |
| PedBE | 0.096 [-0.08, 0.27] | 0.087 | 1.097 | 0.274 | N.S. |
| TeloLength | -0.286 [-0.39, -0.18] | 0.052 | -5.514 | $5.5 \times 10^{-08}$ | *** |

Table S7b: Summary of Regression Results for High YSR Total (Females)

| Model | β estimate [95% CI] | Std Error | t statistic | p | sig |
|---|---|---|---|---|---|
| Phase-Specific_A2 | 0.707 [-0.08, 1.49] | 0.386 | 1.833 | 0.076 | . |
| ENIGMA | 5.316 [0.40, 10.23] | 2.419 | 2.197 | 0.035 | * |
| Centile | 2.945 [0.56, 5.33] | 1.175 | 2.508 | 0.017 | * |
| DevBrainAge | 0.128 [-1.12, 1.37] | 0.612 | 0.209 | 0.835 | N.S. |
| PyBrainAge | -0.039 [-4.79, 4.71] | 2.337 | -0.017 | 0.987 | N.S. |

Note: DNA-based biological age measures not available at this time-point (Y13)

Table S7c: Summary of Regression Results for High YSR Total (Males)

| Model | β estimate [95% CI] | Std Error | t statistic | p | sig |
|---|---|---|---|---|---|
| Phase-Specific_A2 | -0.853 [-1.57, -0.14] | 0.354 | -2.412 | 0.020 | * |
| ENIGMA | -1.113 [-4.53, 2.30] | 1.691 | -0.658 | 0.514 | N.S. |
| Centile | -0.857 [-2.92, 1.21] | 1.023 | -0.838 | 0.407 | N.S. |
| DevBrainAge | -0.110 [-1.01, 0.79] | 0.443 | -0.248 | 0.806 | N.S. |
| PyBrainAge | -3.706 [-9.29, 1.88] | 2.766 | -1.340 | 0.188 | N.S. |

Note: DNA-based biological age measures not available at this time-point (Y13)

Table S7d: Summary of Regression Results for Cumulative ELA

| Model | β estimate [95% CI] | Std Error | t statistic | p | sig |
|---|---|---|---|---|---|
| Phase-Specific_A2 | -0.083 [-0.16, -0.01] | 0.038 | -2.178 | 0.032 | * |
| ENIGMA | -0.375 [-1.38, 0.63] | 0.505 | -0.743 | 0.460 | N.S. |
| Centile | -0.427 [-0.79, -0.06] | 0.183 | -2.338 | 0.022 | * |
| DevBrainAge | -0.012 [-0.18, 0.16] | 0.086 | -0.141 | 0.888 | N.S. |
| PyBrainAge | 0.131 [-0.59, 0.85] | 0.362 | 0.362 | 0.718 | N.S. |
| PedBE | 0.041 [-0.11, 0.19] | 0.075 | 0.550 | 0.583 | N.S. |
| TeloLength | -0.048 [-0.10, 0.00] | 0.027 | -1.800 | 0.073 | . |

Table S7e: Summary of Regression Results for High PSS (Females)

| Model | β estimate [95% CI] | Std Error | t statistic | p | sig |
|---|---|---|---|---|---|
| Phase-Specific_A2 | 1.033 [0.16, 1.90] | 0.429 | 2.408 | 0.021 | * |
| ENIGMA | -0.564 [-6.12, 4.99] | 2.739 | -0.206 | 0.838 | N.S. |
| Centile | 0.116 [-2.18, 2.41] | 1.133 | 0.102 | 0.919 | N.S. |
| DevBrainAge | -0.306 [-1.30, 0.68] | 0.489 | -0.626 | 0.535 | N.S. |
| PyBrainAge | -2.423 [-11.44, 6.59] | 4.448 | -0.545 | 0.589 | N.S. |
| PedBE | -0.166 [-0.74, 0.41] | 0.290 | -0.571 | 0.569 | N.S. |
| TeloLength | 0.123 [-0.21, 0.45] | 0.166 | 0.744 | 0.458 | N.S. |

Table S7f: Summary of Regression Results for High PSS (Males)

| Model | β estimate [95% CI] | Std Error | t statistic | p | sig |
|---|---|---|---|---|---|
| Phase-Specific_A2 | -0.655 [-1.46, 0.15] | 0.400 | -1.639 | 0.107 | N.S. |
| ENIGMA | -1.949 [-6.16, 2.26] | 2.097 | -0.930 | 0.357 | N.S. |
| Centile | -1.514 [-3.81, 0.78] | 1.142 | -1.326 | 0.191 | N.S. |
| DevBrainAge | -0.438 [-1.41, 0.53] | 0.484 | -0.905 | 0.370 | N.S. |
| PyBrainAge | -2.187 [-6.96, 2.59] | 2.379 | -0.919 | 0.362 | N.S. |
| PedBE | 0.028 [-0.65, 0.71] | 0.343 | 0.083 | 0.934 | N.S. |
| TeloLength | -0.162 [-0.34, 0.02] | 0.093 | -1.746 | 0.084 | . |

Supplementary Table S8: Model performance metrics across selected A2 BrainAge model with pre-trained BrainAge models for motion subset (n = 638)

| Metric | Sex | BrainAge Estimates | | | | | |
|---|---|---|---|---|---|---|---|
| | | Phase-Specific_A2 | Phase-Specific_A2 (M) | ENIGMA | Centile | DevBrainAge | PyBrainAge |
| Early-Childhood (n = 380 scans) | | | | | | | |
| Median PAD (IQR) | Female | 0.01 (1.37) | -0.05 (1.5) | 12.07 (7.42) | 2.88 (2.3) | 4.24 (1.7) | 5.68 (4.99) |
| MAE (IQR) | Female | 0.68 (0.72) | 0.74 (0.78) | 12.07 (7.41) | 2.93 (2.25) | 4.24 (1.7) | 5.68 (4.99) |
| Age r [95% CI] | Female | 0.58 [0.47, 0.68] | 0.53 [0.41, 0.64] | 0.02 [-0.14, 0.18] | 0.38 [0.24, 0.5] | 0.44 [0.3, 0.56] | 0.27 [0.12, 0.41] |
| Median PAD (IQR) | Male | -0.15 (1.35) | -0.12 (1.22) | 15.59 (6.96) | 2.6 (2.63) | 4.29 (1.69) | 5.75 (4.48) |
| MAE (IQR) | Male | 0.69 (0.74) | 0.62 (0.68) | 15.59 (6.96) | 2.68 (2.46) | 4.29 (1.69) | 5.75 (4.48) |
| Age r [95% CI] | Male | 0.46 [0.35, 0.56] | 0.51 [0.41, 0.6] | 0.03 [-0.1, 0.16] | 0.38 [0.26, 0.49] | 0.33 [0.2, 0.44] | 0.21 [0.08, 0.33] |
| Late-Childhood (n = 258 scans) | | | | | | | |
| Median PAD (IQR) | Female | 0.04 (1.88) | 0.16 (1.67) | 11.9 (8.38) | 3.42 (2.88) | 2.56 (1.97) | 10.33 (8.14) |
| MAE (IQR) | Female | 0.94 (1.06) | 0.86 (1.25) | 11.9 (8.3) | 3.42 (2.87) | 2.56 (1.88) | 10.33 (8.14) |
| Age r [95% CI] | Female | 0.79 [0.71, 0.85] | 0.8 [0.72, 0.85] | 0.4 [0.24, 0.54] | 0.7 [0.59, 0.78] | 0.77 [0.68, 0.83] | 0.49 [0.34, 0.62] |
| Median PAD (IQR) | Male | 0.08 (1.97) | 0.17 (2.01) | 12.75 (6.72) | 2.21 (2.96) | 2.39 (2.02) | 7.51 (8.28) |
| MAE (IQR) | Male | 1.1 (1.09) | 1.01 (1.12) | 12.75 (6.72) | 2.28 (2.58) | 2.39 (2.02) | 7.51 (8.28) |
| Age r [95% CI] | Male | 0.72 [0.64, 0.79] | 0.71 [0.62, 0.78] | 0.41 [0.26, 0.53] | 0.63 [0.52, 0.72] | 0.74 [0.66, 0.81] | 0.54 [0.41, 0.65] |

Note: Phase-Specific_A2 (M), Metrics for Phase-Specific A2 model trained on the subset of scans that passed MRI motion criteria applied to motion subset Test dataset

Supplementary Table S9: Adversity Score Calculation

| **Component** | **Assessment & Criteria** |
|---|---|
| Birthweight centile[a] at extremes | <10, >90 |
| Low gestational age | <= 37 weeks |
| Mother smoking during pregnancy | Self-reported current smoker |
| Low monthly household income | < $2000 SGD |
| Low family function | Family Assessment Device (FAD) general functioning at age 6<br>>= $85^{th}$ percentile (2.1667) |
| Adverse maternal mental health during pregnancy | > $85^{th}$ percentile for any of the following:<br>Edinburgh Postnatal Depression Scale > 11<br>Beck's Depression Inventory > 14<br>State-Trait Anxiety Inventory > 92 |
| Adverse maternal physical health during pregnancy | Long-term illness at pregnancy week 11<br>Gestational diabetes mellitus diagnosis<br>Hypertension diagnosis |

a. Birthweight centile calculated with Intergrowth [25]
Note: Most prenatal assessments conducted at pregnancy week 26 unless stated otherwise
